# A combination of mRNA features influence the efficiency of leaderless mRNA translation initiation

**DOI:** 10.1101/2020.05.06.081141

**Authors:** Mohammed-Husain M. Bharmal, Alisa Gega, Jared M. Schrader

## Abstract

Bacterial translation is thought to initiate by base-pairing of the 16S rRNA and the Shine-Dalgarno sequence in the mRNA’s 5’UTR. However, transcriptomics has revealed that leaderless mRNAs, which completely lack any 5’UTR, are broadly distributed across bacteria and can initiate translation in the absence of the Shine-Dalgarno sequence. To investigate the mechanism of leaderless mRNA translation initiation, synthetic *in vivo* translation reporters were designed that systematically tested the effects of start codon accessibility, leader length, and start codon identity on leaderless mRNA translation initiation. Using this data, a simple computational model was built based on the combinatorial relationship of these mRNA features which can accurately classify leaderless mRNAs and predict the translation initiation efficiency of leaderless mRNAs. Thus, start codon accessibility, leader length, and start codon identity combine to define leaderless mRNA translation initiation in bacteria.

## INTRODUCTION

Translation initiation is a critical step for fidelity of gene expression in which the ribosome initiation complex is formed on the start codon of the mRNA. Since the canonical start codon, AUG, compliments both initiator and elongator methionyl-tRNAs, the ribosome must distinguish the start AUG codon from elongator AUG codons. Incorrect initiation at an elongator AUG can lead to non-functional products that can be detrimental to cellular fitness (1–3). Canonical start codon selection is thought to occur by the base-pairing of the 16S rRNA with a Shine-Dalgarno (SD) sequence in the mRNA located 5nt upstream of the start codon (4–6). The base pairing between the 16S rRNA and mRNA was shown to be critical for initiation since mutation of the anti-SD (aSD) in the 16S rRNA is lethal (7), and translation of a gene lacking a canonical SD sequence could be restored when the 16S of the rRNA were mutated to a complimentary sequence (8). While the SD-aSD pairing clearly impacts translation initiation efficiency (TIE) in *E. coli*, other studies have found that the SD:aSD interaction is not essential for correct selection of the start codon (9,10). Indeed, “orthogonal” ribosomes with altered 16S rRNA aSD sequences were found to initiate at the normal start codons throughout the transcriptome (11). Interestingly, *E. coli* lacks SD sites within its genome in approximately 30% of its translation initiation regions (TIRs) with other species of bacteria containing SD sites in as few as 8% of their TIRs (12,13). Indeed, RNA-seq based transcription mapping experiments have found that many bacterial mRNAs are “leaderless” and begin directly at the AUG start codon (14–16), and that these mRNAs are abundant in pathogens such as *M. tuberculosis* and in the mammalian mitochondria (17).

To account for the lack of essentiality of the SD site, a “Unique accessibility model” was proposed which posited that start codon selection occurs due to the TIR being accessible to initiating ribosomes, while elongator AUGs are physically inaccessible due to RNA secondary structures (18). This model was based upon a strong negative correlation observed between mRNA secondary structure content in the TIR and TIE (19–21). This model is further supported by genomic analysis of RNA secondary structure prediction of mRNA TIRs in which there’s a lower amount of secondary structure in the TIR compared to elongator regions, which is conserved across all domains of life (22). While the unique accessibility model is overly simplistic, more advanced computational approaches have been able to combine TIR accessibility with SD strength, spacing, and standby sites to more accurately predict TIE of leadered mRNAs (23). While TIR accessibility has been shown to be critical in many leadered mRNAs, it has not yet been systematically tested for leaderless mRNAs.

Genome-wide RNA-seq transcript mapping experiments have revealed that leaderless mRNAs are widespread across bacteria (14), yet little is known about their mechanism of translation initiation. While very few leaderless mRNAs has been identified in *E. coli* (0.7% leaderless mRNAs (24)), other bacteria and archaea contain a large majority of their transcripts as leaderless mRNAs (up to 72% leaderless mRNAs (14,25)). Additionally, sizeable proportions of leaderless mRNAs have been identified in bacteria of clinical significance, such as *Mycobacterium tuberculosis*, and of industrial significance like *Corynebacterium glutamicum* (15,26). In the model bacterium *Caulobacter crescentus* approximately 17% of mRNAs are leaderless (27), with the fastest doubling time known of any bacterium with large numbers of leaderless mRNAs. In addition, *C. crescentus* has good genetic tools, making it an ideal model to study translation initiation of leaderless mRNAs.

Importantly, the role of TIR accessibility has not been systematically tested for leaderless mRNAs, however, some aspects of their initiation have been identified which are distinct from leadered mRNAs. Mitochondrial leaderless mRNAs have been found to lack 5’ secondary structure (28), in support of a TIR accessibility model. Additionally, mutagenesis of the *Mycobacterium smegmatis pafA* leaderless mRNA to perturb its secondary structure showed that secondary structure content negatively correlated with this translation levels (29). However, the changes in codon usage across the mutants make the relative impact of secondary structure and codon usage unknown for this mRNA. In opposition to the canonical initiation mechanism, leaderless mRNAs can initiate with 70S ribosomes where IF2 is known to stimulate their translation, and IF3 can inhibit leaderless translation (30,31). Additionally, AUG is the most efficient start codon in leaderless mRNAs in *E. coli* or *Haloarchaea*, (32–36), while AUG or GUG are both efficient leaderless mRNA start codons in *M. smegmatis* (16). In *E. coli*, suppressor tRNAs could restore initiation on non-AUG codons for leadered RNAs, but not for leaderless RNAs (32), suggesting that for leaderless mRNAs an AUG start codon has unique initiation properties independent of perfect codon-anticodon base-pairing. Indeed, genomic prediction of leaderless mRNAs suggests a very high preference of AUG (79%) at the 5’ end of leaderless mRNAs; with a smaller percentage of GUG (10%), UUG (6%) and others (3%) (13). In addition to the start codon identity, TIE of mRNAs with short leaders (<5nt) is significantly lower as compared to their fully leaderless counterparts (34,35,37–39). Altogether, this suggests that leaderless mRNAs strongly prefer AUG and are inhibited by having short leaders.

In order to understand the mRNA sequence features needed for leaderless translation initiation, we systematically measured the effect of TIR accessibility, start codon identity, and leader length on leaderless mRNA translation initiation in *C. crescentus*. Using synthetic *in vivo* translation initiation reporters, we show that TIR accessibility, start codon identity, and leader length all dramatically affect leaderless mRNA TIE. The dependencies of each mRNA feature on TIE were then built into a simple computational model (TIE_leaderless_ model) that accurately predicts which RNAs in the *C. crescentus* transcriptome would be initiated as leaderless RNAs with an area under the curve (A.U.C.) of a Receiver Operator Characteristic (ROC) curve of 0.99. The TIE_leaderless_ model also accurately predicts the translation initiation efficiency of *in vivo* leaderless mRNA reporters (R^2^=0.87). This therefore provides the first systematic analysis of mRNA features required for leaderless initiation and the *C. crescentus* TIE_leaderless_ model will likely provide a foundation for our understanding of leaderless mRNA translation initiation across bacteria.

## MATERIAL AND METHODS

### Computational predictions of start codon accessibility

#### Retrieving transcript sequences

All the RNA sequences were retrieved from transcription start sites and translation start site data available from RNA-seq and ribosome profiling respectively (27,40) using the *C. crescentus* NA1000 genome sequence (41). For *M. smegmatis*, RNA-seq and ribosome profiling data were downloaded from the European Nucleotide Archive (https://www.ebi.ac.uk/arrayexpress/experiments/E-MTAB-2929/) and for *M. tuberculosis*, RNA-seq data was obtained from Gene Expression Omnibus (GEO) accession number GSE62152 and analyzed using the CP000480.1 and NC_000962 genome sequences respectively (16). For *H. volcanii*, RNA-seq and ribosome profiling were provided by The DiRuggerio lab, and analyzed with the *H. volcanii* NCBI RefSeq genome (taxonomy identification [taxid] 2246; 1 chromosome, 4 plasmids) (42). For *<u>M. musculus</u>* mitochondria, RNA-seq and ribosome profiling data were downloaded from (43) and analyzed with the NC_005089 genome sequence. The TIR sequences were then extracted from all open reading frames (ORFs) using 50 nt (25 nt upstream of start codon and 25 nt downstream from start codon). If the 5’ upstream untranslated region (UTR) was less than 25 nt, then 50 nt from transcription start site was used for all TIR calculations. Classification of mRNA type (leaderless, non-SD, or SD) were obtained from Schrader *et al*. PLOS Genetics 2014.

#### Translation Efficiency Data

Ribosome profiling and RNA-seq Translation efficiency data were obtained for *C. crescentus* from (27). Ribosome profiling and RNA-seq Translation efficiency data were obtained for *H. volcanii* from the group of Prof. Jocelyn DiRuggerio (44). Ribosome profiling and RNA-seq sequencing data were obtained for *M. smegmatis* from the European Nucleotide Archive (https://www.ebi.ac.uk/arrayexpress/experiments/E-MTAB-2929/)(16). For both ribosome profiling and RNA-seq data the sequencing reads were downloaded as fastq files and the adapter poly-A sequences were trimmed using a custom python script. Trimmed reads were then depleted of rRNA and tRNA reads by alignment with bowtie (45), and the remaining non-rRNA/tRNA reads were then aligned to the *M. smegmatis* MC2 155 genome. RPKM values were then calculated based upon the CP009494.1 annotation. To avoid confounding effects from initiating or terminating ribosomes, the first 15nt and last 15nt of ORFs were omitted from the RPKM calculations. ORFs with less than 50 reads in a given sample were omitted from the RPKM calculation. The Translation Efficiency (TE) was then calculated as the ratio of the RPKM_Ribosome profiling_/RPKM_RNA-seq_.

#### Calculation of ΔG_unfold_

Start codon accessibility was computed similar to (46) by comparing the native TIR RNA structure (ΔG_mRNA_) to that of the same TIR bound by an initiating ribosome (ΔG_init_). Since ribosome binding requires a single-stranded region of the mRNA we approximated this by forcing the TIR to be single stranded. The overall calculation was performed in three steps:

1. Calculation of ΔG_mRNA_: The minimum free energy (mfe) labelled as ΔG_mRNA_ was calculated using RNAfold web server of the Vienna RNA websuite (47) at the growth temperature of each organism by inputting all the TIR sequences in a text file using command line function ‘RNAfold --temp “temp” <input_sequences.txt >output.txt’. The output file was in the default RNAfold format with each new sequence on one line followed by dot-bracket notation (Vienna format) in the next line. RNAstructure (48) was used to generate ct files for each of the mfe structures predicted in RNAfold which contained all the base pair indexes for each sequence.
2. Calculation of ΔG_init_: The base pairs in the TIR (from up to 12 nt upstream of the start codon to 13 nt downstream of the start codon) were broken and forced to be single stranded including any pairs formed from the TIR and outside. If the 5’UTR length was more than or equal to 25 nt, then the RBS was selected from −12 to +13 nt (25 nt). If the 5’UTR length was less than 25, then the TIR comprised of the entire 5’UTR to +13 nt. A new dot bracket file with these base-pairing constraints was then used in the RNAfold program (47) with the same RNA sequence to calculate the ΔG_init_.
3. Calculation of ΔG_unfold_: Lastly, ΔG_unfold_ was calculated by subtracting ΔG_mRNA_ (mfe of mRNA in native state) from ΔG_init_ (mfe of mRNA after ribosome binding) (eq 1. ΔG_unfold =_ ΔG_init_ − ΔG_mRNA_).

### Cell growth and media

#### *E. coli* culture

For cloning, plasmids with the reporter gene were transformed in *E. coli* top10 competent cells using heat shock method for 50-55 secs at 42°C. Luria-Bertani (LB) liquid media was used for outgrowth and the colonies were plated on LB/kanamycin (50 μg/mL) agar plates. For miniprep, the *E. coli* cultures were inoculated overnight(O/N) in liquid LB/kanamycin (30 μg/mL).

#### *C. crescentus* culture

For cloning, plasmids were transformed in NA1000 *C. crescentus* cells after sequence verification using electroporation. The *C. crescentus* NA1000 cells were grown in Peptone Yeast Extract (PYE) liquid medium. After transformation, for the outgrowth liquid PYE medium was used (2mL) and then plated on PYE/kanamycin (25 μg/mL) agar plates. For imaging, the *C. crescentus* culture were grown O/N at different dilutions in liquid PYE/kanamycin (5 μg/mL). Next day, the cultures growing in log phase were diluted and induced in liquid PYE with kanamycin (5 μg/mL) and Xylose (final concentration of 0.2%) such that the optical density (OD) was around 0.05 to 0.1.

### Design and generation of translation reporters

#### Oligos and plasmid design

For the design and generation of reporter assay, a plasmid with a reporter gene (yellow fluorescent protein (YFP)), under the control of an inducible xylose promoter was used. The pBYFPC-2 plasmid containing the kanamycin resistant gene was originally generated from (49). A list of oligos used for generating plasmids with different 5’ UTRs of YFP is attached as a supplementary table (Table S6).

#### Inverse PCR mutagenesis and Ligation

The 5’UTR region and start codon of the YFP reporter protein was replaced with other TIR sequences. This was done by inverse PCR, in which the leaderless TIR is attached to the reverse primer as an overhang. Initial denaturation was done at 98°C for 5 mins. Followed by 30 cycles of denaturation at 98°C for 10 secs, annealing at 60°C for 10 secs and extension at 72°C for 7 mins and 20 secs. After 30 cycles, final extension was done at 72°C for 5 mins. The polymerase used was Phusion (Thermoscientific 2 U/μL). The PCR product was then DPNI treated to cut the template DNA using DPNI enzyme (Thermoscientific 10 U/μL). The DPNI treated sample was then purified using Thermo fisher GeneJET PCR Purification kit. The purified sample (50 ng) was then used for blunt end ligation using T4 DNA Ligase (Thermoscientific 1 WeissU/μL).

#### Transcription reporter design

For the design of transcription reporter assay, a plasmid with a 28 nt mutant version of 5’ UTR (CCNA_03971) in front of reporter gene (yellow fluorescent protein (YFP)) was used. This reporter gene had the nucleotide A at its +1 position and the reporter gene was under the control of an inducible xylose promoter. The pBYFPC-2 plasmid containing the kanamycin resistant gene was originally generated from (49). The +1 nucleotide was mutated to all other nucleotides (G, C or T) and these 3 mutant plasmids were synthesized into DNA oligos and cloned by Genscript.

The insertion sequence from the +1 nt (underlined) to 28 nt including start codon (atg) to the RBS for each construct is shown below:

~~~
A.) <u>**a**</u>ccgattaacg**atg**gtggttgttctggc
C.) <u>**c**</u>ccgattaacg**atg**gtggttgttctggc
G.) <u>**g**</u>ccgattaacg**atg**gtggttgttctggc
T.) <u>**t**</u>ccgattaacg**atg**gtggttgttctggc
~~~

#### Transformation in *E. coli* cells

5 μL of the ligation reaction was then added to 50 μL of *E. coli* top10 competent cells. Then the mixture was incubated in ice for 30 mins. Then heat shocked for 50-55 secs in the water bath at 42°C. Then immediately kept in ice for 5 mins, after which 750 μL of LB liquid medium was added to the cells for outgrowth and kept for incubation at 37°C for 1 hr at 200 rpm. After this, 200-250 μL of the culture was plated on LB/kanamycin (50 μg/mL) agar plates.

#### Colony screening and sequence verification

The colonies grown on LB/kanamycin plates were screened by colony PCR to first screen for the presence of the new TIR insert. The cloning results in the replacement of the larger 5’UTR region of YFP with a smaller region containing a leaderless TIR, thus distinguished easily on an analytical gel. The forward and reverse primer used for the screening results in approximately 180 base pairs, whereas the original fragment amplified with the same oligos is 245 base pairs. The forward oligo used was pxyl-for: cccacatgttagcgctaccaagtgc and reverse oligo is eGYC1: gtttacgtcgccgtccagctcgac. Upon verification, a small aliquot (4 μL) of the colony saved in Taq polymerase buffer was inoculated in 5 mL of liquid LB/kanamycin (30 μg/mL) and incubated overnight at 37°C at 200 rpm. Next day, the culture was miniprepped using Thermo fisher GeneJET Plasmid Miniprep kit. The concentration of DNA in the miniprepped samples were measured using Nanodrop 2000C from Thermoscientific. DNA samples were sent to Genewiz for sanger sequencing to verify the correct insert DNA sequences using the DNA primer eGYC1: gtttacgtcgccgtccagctcgac (49).

#### Transformation in *C. crescentus* NA1000 cells

After the sequences were verified, the plasmids were transformed in *C. crescentus* NA1000 cells. For transformation, the NA1000 cells were grown overnight at 28°C in PYE liquid medium at 200rpm. The next day, 5 mL of cells were harvested for each transformation, centrifuged and washed three times with autoclaved milliQ water. Then, 1 μL of sequence verified plasmid DNA was mixed with the cells and electroporated using Bio-Rad Micropulser (program Ec1 set at voltage of 1.8 kV). Then, the electroporated cells were immediately inoculated in 2 mL of PYE for 3 hours at 28°C at 200rpm. Then 10-20 μL of culture was plated on PYE/ kanamycin agar plates. Kanamycin-resistant colonies were grown in PYE/kanamycin media overnight and then stored as a freezer stock in the - 80°C freezer

### Cellular assay of translation reporters

*C. crescentus* cells harboring reporter plasmids were serially diluted and grown overnight in liquid PYE/kanamycin medium (5 μg/mL). The next day, cells in the log phase were diluted with fresh liquid PYE/kanamycin (5 μg/mL) to have an optical density (OD) of 0.05-0.1. The inducer xylose was then added in the medium such that the final concentration of xylose is 0.2%. The cells were grown for 6 hours at 28°C at 200 rpm. After this, 2-5 μL of the cultures were spotted on M2G 1.5% agarose pads on a glass slide. After the spots soaked into the pad, a coverslip was placed on the pads and the YFP level was measured using fluorescence microscopy using a Nikon eclipse NI-E with CoolSNAP MYO-CCD camera and 100x Oil CFI Plan Fluor (Nikon) objective. Image was captured using Nikon elements software with a YFP filter cube with exposure times of 30ms for phase-contrast images and 300 ms for YFP images respectively. The images were then analyzed using a plugin of software ImageJ (50) called MicrobeJ (51).

### Three component model calculations and leader length/identity analysis

For all RNA transcripts in the *C. crescentus* genome identified in (27,40), we computed their capacity to initiate as a leaderless mRNA using equation 2: (TIE_Leaderless mRNA*(k)*_ = Max TIE (1) - (1-TIE_ΔGunfold_) - (1-TIE_start codon identity(j)_) − (1-TIE_leader length(i)_) where k= a given RNA transcript, j=start codon identity, and i=leader length(nt). To identify putative leaderless mRNA TIRs, we first asked if the 5’ end contained an AUG or near cognate start codon, and if not we scanned successively from the 5’ end for AUG trinucleotides within the first 8 nt. Near cognate start codons were omitted from positions containing leader nucleotides since AUG codons yielded higher TIE values even in the presence of a leader. We next asked if there is an AUG or near cognate start codon further downstream by scanning 5’ to 3’ through the first 18 nt. If found, we calculated TIE_leaderless mRNA_ with all different possible cognate/near-cognate start codons along the TIR. Then of all the different possibilities, the one having the highest TIE_leaderless_ score was selected for further analysis (Fig 7A).

To utilize TIE_leaderless mRNA_ for classification, each RNA was then categorized into two different classes based on 5’ end sequencing data and ribosome profiling based global assays ((27,40)): true leaderless – RNAs that are known to initiate directly at a 5’ start codon (judged by a complete lack of a 5’ UTR and a ribosome density >1/20 the downstream CDS (27)), and false leaderless – RNAs that are not initiated at a 5’ start codon. A small subset was classified as “unknown”, as they contain very short leaders and lack SD sites, making their mode of translation initiation ambiguous. Using these TIE_leaderless mRNA_ values, a ROC curve was plotted using scikit-learn library in python (52) with the “true leaderless” and “false leaderless” RNAs (TIE_leaderless mRNA_ values for the *C. crescentus* transcriptome can be found in Table S1).

To utilize TIE_leaderless mRNA_ for prediction of translation initiation reporter levels, we first converted all negative TIE_leaderless mRNA_ scores to zero. Next, we compared the TIE_leaderless mRNA_ scores to the YFP levels of the translation initiation and performed a linear regression calculation using the linest function in microsoft excel and libreoffice calc. For prediction of native leaderless mRNA translation levels, TE measurements from ribosome profiling experiments (27) were compared to the TIE_leaderless mRNA_ scores.

## RESULTS

### Computational prediction of C. crescentus start codon accessibility

To assess the role of mRNA accessibility across mRNA types, ΔG_unfold_ calculations were performed on all *C. crescentus* translation initiation regions (TIRs). ΔG_unfold_ represents the amount of energy required by the ribosome to unfold the mRNA at the translation initiation region (TIR) and has been identified as a metric that correlates with translation efficiency in *E. coli* (46). ΔG_unfold_ was calculated for all TIRs by first predicting the minimum free energy of the 50 nt region of the mRNA (ΔG_mRNA_) around the start codon using RNAfold (47). ΔG_init_ was then calculated in which the TIR (25nt surround the start codon), roughly equivalent to a ribosome footprint, was constrained to be single stranded to approximate accessibility for the ribosome to initiation. ΔG_unfold_ was then calculated using equation 1 (eq 1. ΔG_unfold =_ ΔG_init_ − ΔG_mRNA_) which represents the energy required to open the TIR to facilitate translation initiation (Fig 1A). ΔG_unfold_ calculations were performed on all the CDSs in the genome (Fig 1B) and classified into mRNA types based on transcriptome and ribosome profiling maps of the *C. crescentus* genome (27). The transcripts were categorized into two major classes: leaderless (no 5’ UTR) and leadered (those containing a 5’ UTR). Leadered mRNAs were further categorized into subclasses based upon the presence of the Shine-Dalgarno (SD) sequence (27). Shine-Dalgarno (SD) (containing a SD sequence in the 5’ UTR) and nonSD (lacking an SD sequence in the 5’ UTR). Since it is also known that some polycistronic operons reinitiate translation between CDSs without dissociation of the ribosomal subunits, we also examined the ΔG_unfold_ of TIRs occurring downstream of the first CDS in polycistronic mRNAs (Operons). The average ΔG_unfold_ value of leaderless mRNAs (5.6 kcal/mol) was significantly lower than SD (11.9 kcal/mol, p= 1.5E-105), nonSD (10.3 kcal/mol, p= 6.9E-71) and internal operon TIRs (13.2 kcal/mol,p= 1.1E-143) as calculated by pairwise 2-sided T-tests with unequal variance (Fig 1B). The lower ΔG_unfold_ values of nonSD TIRs may be due to the loss of stabilization of TIRs from base pairing between the anti-SD site in the 16S rRNA and the SD site in the mRNA. We also observed that average ΔG_unfold_ of nonSD TIRs was significantly lower than SD TIRs (p= 1.8E-14) and operon TIRs (p=1.4E-44). The difference between the average ΔG_unfold_ of SD and operon genes was also significant (p=2.1E-09). Because the ribosome is an efficient RNA helicase, it is possible that the increased ΔG_unfold_ of operon TIRs may be tolerated by the ribosome’s ability to unwind such structures when terminating on the previous CDSs. We hypothesized that the low ΔG_unfold_ observed for leaderless mRNAs was due to an intrinsic requirement for their initiation, however, because the size of the leaderless mRNA footprint is significantly smaller than a leadered mRNA footprint, the low ΔG_unfold_ observed for leaderless mRNAs could potentially be explained by the smaller ribosome footprint size. To explore this possibility, we analyzed the ΔG_unfold_ of leadered mRNA TIRs using the same footprint size and region as leaderless mRNAs (13nt) in the ΔG_unfold_ calculation (Fig S1). We observed that the ΔG_unfold_ was still significantly lower for leaderless mRNA TIRs, suggesting that the low ΔG_unfold_ for leaderless mRNA TIRs is not simply an artifact of the smaller mRNA footprint size.

**Figure 1.**
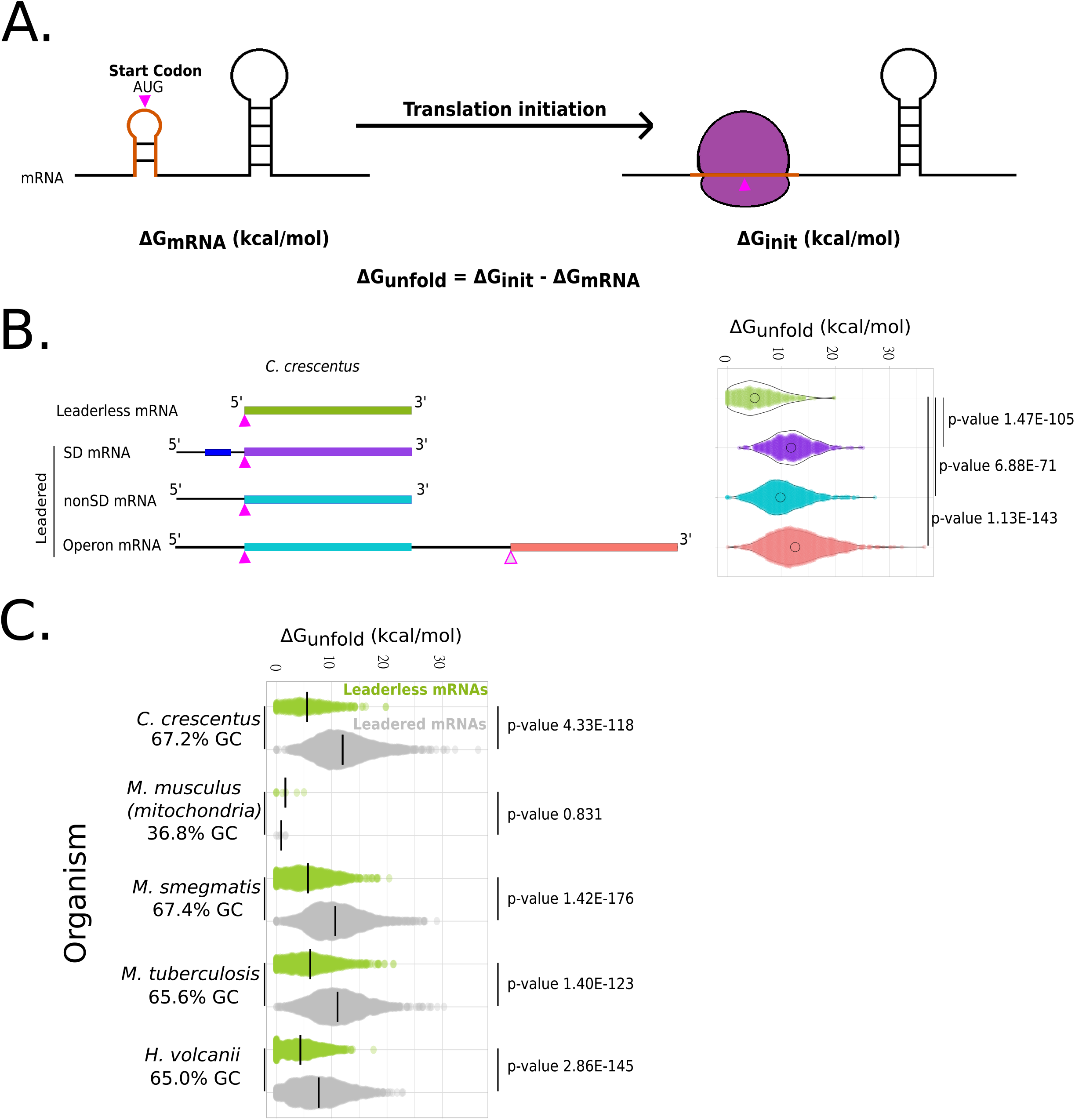
Leaderless mRNA translation initiation regions are more accessible than leadered mRNAs. A.) Predicted unfolding energy of mRNAs. The predicted mRNA minimum free energy (ΔG_mRNA_) is represented on the left. The orange translation initiation region indicates a ribosome footprint surrounding the start codon (pink). The image on the right represents the mRNA upon initiation (ΔG_init_) where the orange initiation region is unfolded. The ΔG_unfold_ represents the amount of energy required by the ribosome to unfold the translation initiation region of the mRNA. B.) Violin plots of ΔG_unfold_ (right) calculated for all the mRNAs of each class (left) in the *Caulobacter crescentus* genome based on the transcript architecture(27,40). P-values were calculated based on t-test (two tailed, unequal variance).,,

To explore whether low ΔG_unfold_ for leaderless mRNA TIRs a species-specific property of *C. crescentus*, or a general property of leaderless mRNA TIRs, we calculated ΔG_unfold_ for TIRs in other organisms identified to contain a significant number of leaderless mRNAs. We identified two additional bacteria (*Mycobacterium smegmatis* and *Mycobacterium tuberculosis*) (16), one archaeal species (*Haloferax volcanii*) (25,44), and one mitochondrial genome (*Mus musculus*) (43) which had transcriptome information and ribosome profiling or mass spec data supporting a significant number of leaderless mRNAs. Across bacteria and archaea, the leaderless mRNA ΔG_unfold_ remained quite low as compared to their leadered mRNAs counterparts (leaderless average 5.6 to 7.6 kcal/mol, leadered average 12.0-12.9 kcal/mol) (Fig 1C). In *M. musculus* mitochondria however, leaderless mRNAs and leadered mRNAs were both observed to have low ΔG_unfold_ (Fig 1C), perhaps in part due to the relatively low GC%. Since leaderless mRNAs showed a rather low ΔG_unfold_, and lack complexities associated with leadered mRNAs, such as SD or standby sites which are important for leadered initiation (23), we further explored the functional role of ΔG_unfold_ in *C. crescentus* leaderless mRNAs.

### Systematic analysis of C. crescentus leaderless mRNA TIR determinants using in vivo translation reporters

Leaderless mRNAs initiation is known to be strongly influenced by addition of nucleotides prior to the start codon (leader nts) and by start codon identity (32–39); however, the role of TIR accessibility has been poorly described in this class of mRNAs. To understand the role of these three mRNA features we systematically tested each feature using *in vivo* leaderless mRNA translation initiation reporters. Translation initiation reporters were designed in which the start codon of plasmid pBXYFPC-2 was replaced with an AUG fused directly to the +1 nt of the xylose promoter(49). The xylose promoter was chosen because it is one of the best characterized promoters in *Caulobacter* and its TSS was mapped to the same nt by two independent methods(40). An additional 15-24 nt after the 5’ AUG was added to allow complete replacement of the 5’ leader and start codon in pBXYFPC-2 with a leaderless TIR. Since only the first 6-9 codons are altered across leaderless mRNA mutants, and the vast majority of the YFP CDS is unaltered, this allows a sensitive system to measure changes in translation initiation. As leaderless TIR mutants may also alter the amino acid sequence, additional care was also taken to ensure that mutations would not alter the N-end rule amino acid preferences of the resulting proteins (53). Using this *in vivo* translation initiation system, we generated three different sets of leaderless TIR reporters to test the effect of ΔG_unfold_, start codon identity, and additional leader length on *C. crescentus* translation initiation.

As leaderless mRNAs were predicted to have TIRs with low ΔG_unfold_ values, we engineered several RNA hairpins in the TIR to assess the role of ΔG_unfold_ on translation initiation (Fig 2A). Since very few natural *C. crescentus* mRNAs contained RNA structure content in their TIRs (Fig 1B), six synthetic hairpins were designed, varying in stem and loop sizes (Table S2). Into each construct, we also introduced synonymous codon mutations designed to alter the secondary structure content, yielding a range of ΔG_unfold_ values without altering the amino acid sequence within a given hairpin (Table S2). Importantly, the entire range of ΔG_unfold_ values across the synthetic hairpins spans the entire range calculated for natural leaderless mRNAs (Fig 1, Table S2). For all hairpins, we observed that lowering ΔG_unfold_ and thereby increasing the accessibility of the start AUG led to an increase in the level of YFP production (Fig 2B). Since, 6/7 of the hairpin mutant sets showed a relationship in which hairpin codon usage frequency positively correlated with ΔG_unfold_ (Table S2), it is most likely that the observed reduction in YFP reporter levels is a result of increased structure content and is not likely to be caused by faster elongation of common codons in the TIR. Additionally, across all mutant hairpins sets generated, we observed a strong negative correlation between the YFP reporter level and the ΔG_unfold_ across a vast range of values with a linear correlation R^2^ value of 0.84 (Fig 2B). These data suggest that accessibility of the start codon is a critical feature for leaderless mRNA translation initiation.

**Figure 2.**
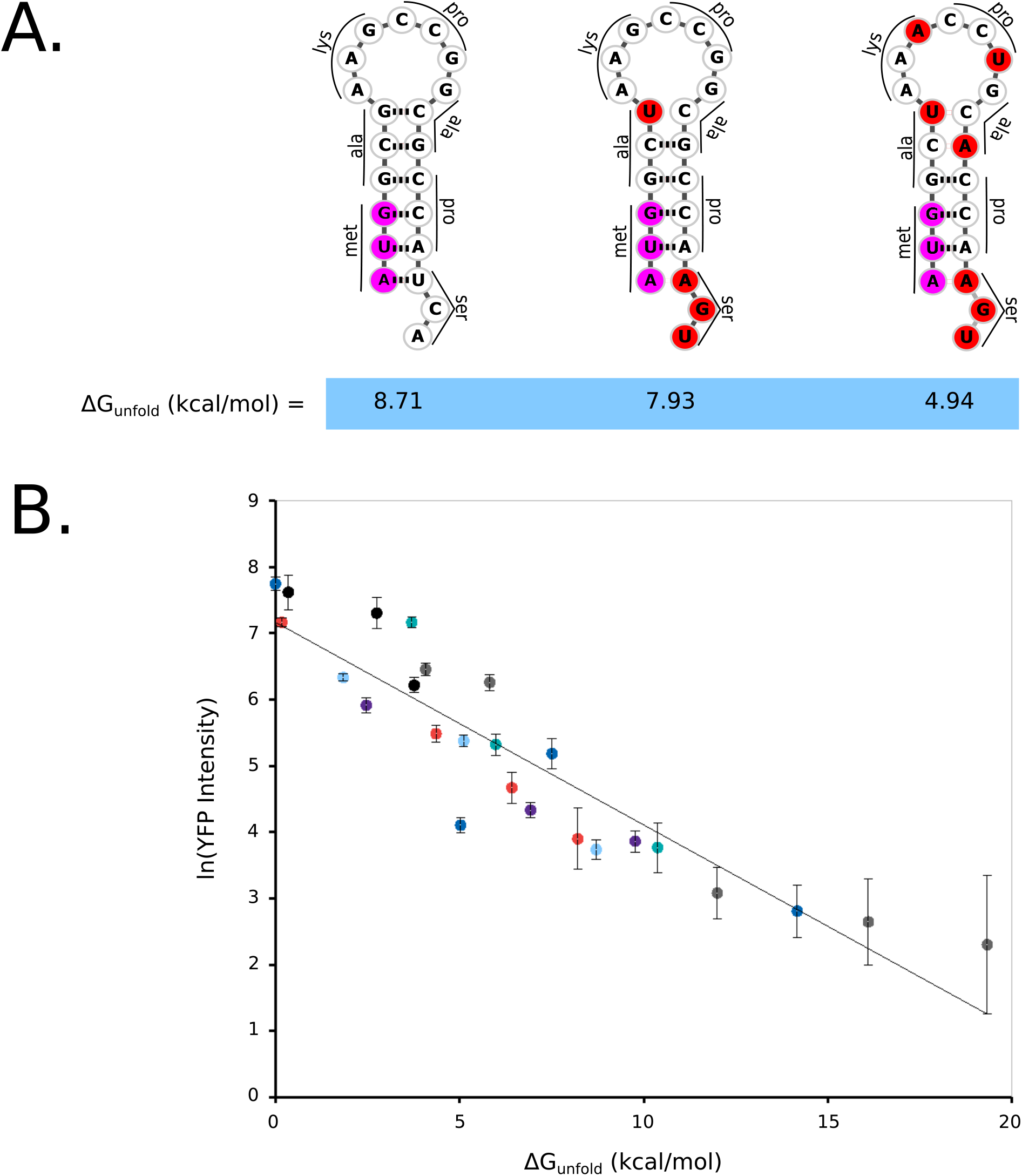
ΔG_unfold_ strongly influences leaderless mRNA translation. A.) A representative TIR synthetic stem loop synonymous mutation set with varying ΔG_unfold_ values. The bases in the start codon are colored pink, red bases highlight where mutations were introduced to disrupt base pairing. B.) *In vivo* translation reporter levels the various leaderless RNA mutants. Each hairpin and its synonymous codon mutant set are shown with the same color (Raw data can be found in Table S1). Black points = leaderless set 1, grey points = leaderless set 2, dark blue points = leaderless set 3, purple points = leaderless set 4, light blue points = leaderless set 5, red points = leaderless set 6, and teal points = leaderless set 7. The natural log of the average YFP intensity per cell is shown and error bars represent the standard deviation of three biological replicates. The dotted blue line represents a linear curve fit with an R^2^ value of 0.84 and a slope of −0.3.

Next, we systematically tested the effect of the start codon identity on the *in vivo* translation initiation reporters. In *C. crescentus*,natural leaderless mRNAs initiate with an AUG, GUG, or UUG start codon (27,40). Since it is well established that start codon identity can affect leaderless mRNA translation initiation (32–36) we generated variants with different start codon identities. Here, AUG was mutated to other near cognate start codons GUG, CUG, UUG, AUC, AUU, AUA which are known to be the start codons of other leadered mRNAs in *C. crescentus* (27). We also included a non-cognate GGG codon as a negative control since no GGG start codons are known to occur in *C. crescentus*.The results showed that replacing the original AUG codon with any of the other near cognate codons drastically decreased the translation initiation reporter levels, while the GGG codon yielded the lowest translation initiation reporter levels (Fig 3). To examine whether the mutation in +1 nt resulted from lower transcription or from lower translation, we generated leadered mRNA reporters with all 4 possible +1 nts and tested their *in vivo* reporter levels as similarly has been performed in *M. smegmatis* (16). A +1 G led to a mild reduction in reporter activity compared to a +1 A (Fig S5), suggesting that most of the observed changes in the leaderless mRNA reporter levels likely come from translation. These data show that the AUG triplet is by far the preferred start codon for *C. crescentus* leaderless mRNAs.

**Figure 3.**
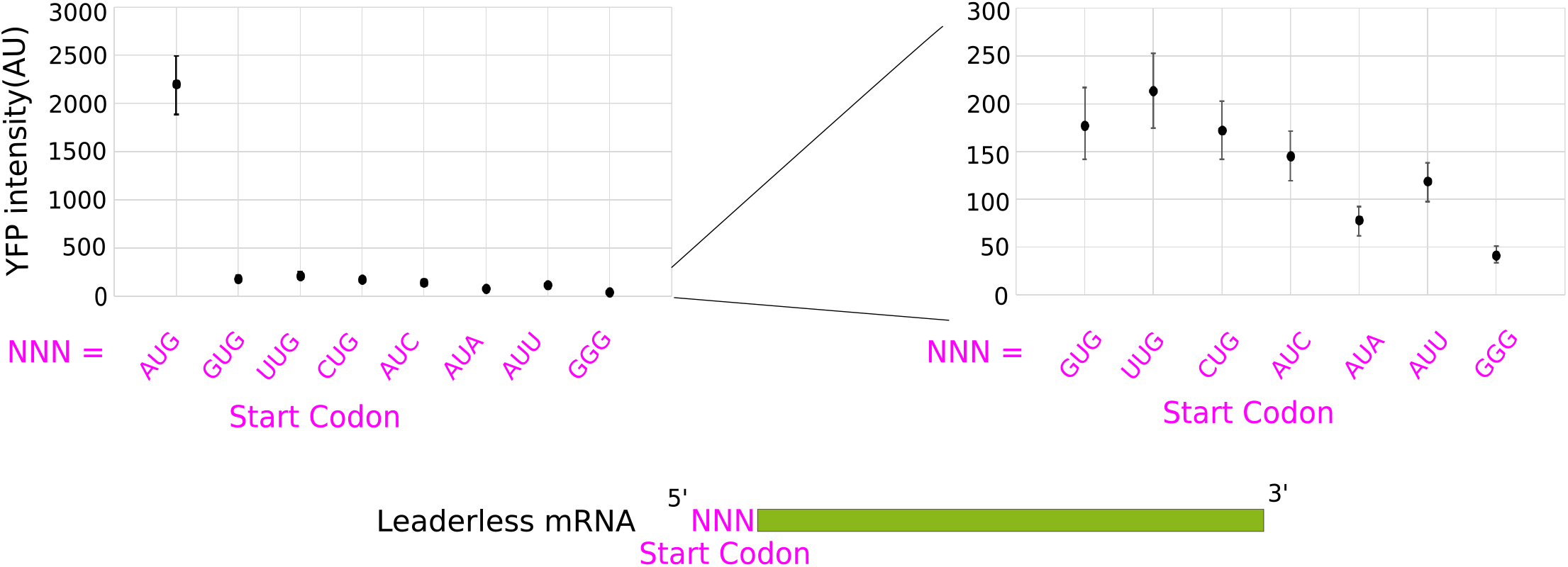
Leaderless mRNAs have a strong preference for AUG start codons. Leaderless mRNA *in vivo* translation reporters were generated with the start codons listed on the X-axis and their average YFP intensity per cell were measured. On the right, is a zoomed in view of all non-AUG codons tested. Error bars represent the standard deviation from three biological replicates.

Finally, we systematically tested the role of additional leader length on *C. crescentus* leaderless mRNAs. In *E. coli*, even a single nucleotide before the AUG is known to inhibit initiation of leaderless mRNAs (37). To test if *C. crescentus* leaderless mRNAs were negatively impacted by leader nucleotides we generated a set of reporters with 0, 1, 2, 3, 5, 10, or 20 5’ Adenosines before the AUG start codon (Fig 4). An A-rich sequence was chosen as it lacks any possible SD sites and is unlikely to form secondary structure, and ΔG_unfold_ values were not altered upon addition of these 5’ bases to the leaderless translation initiation reporter (Table S1). Across this set of mutants, additional nucleotides showed a strong decrease in translation initiation reporter levels with increasing leader length (Fig 4). The translation initiation reporter levels dropped by approximately 2-fold for each additional A that was added to the 5’end (TIE_leader length_ = 0.45×i^-0.91^, R^2^=0.92, i=leader length (nt)). This confirms that even a short leader can lead to a significant reduction in translation initiation of *C. crescentus* leaderless mRNAs.

**Figure 4.**
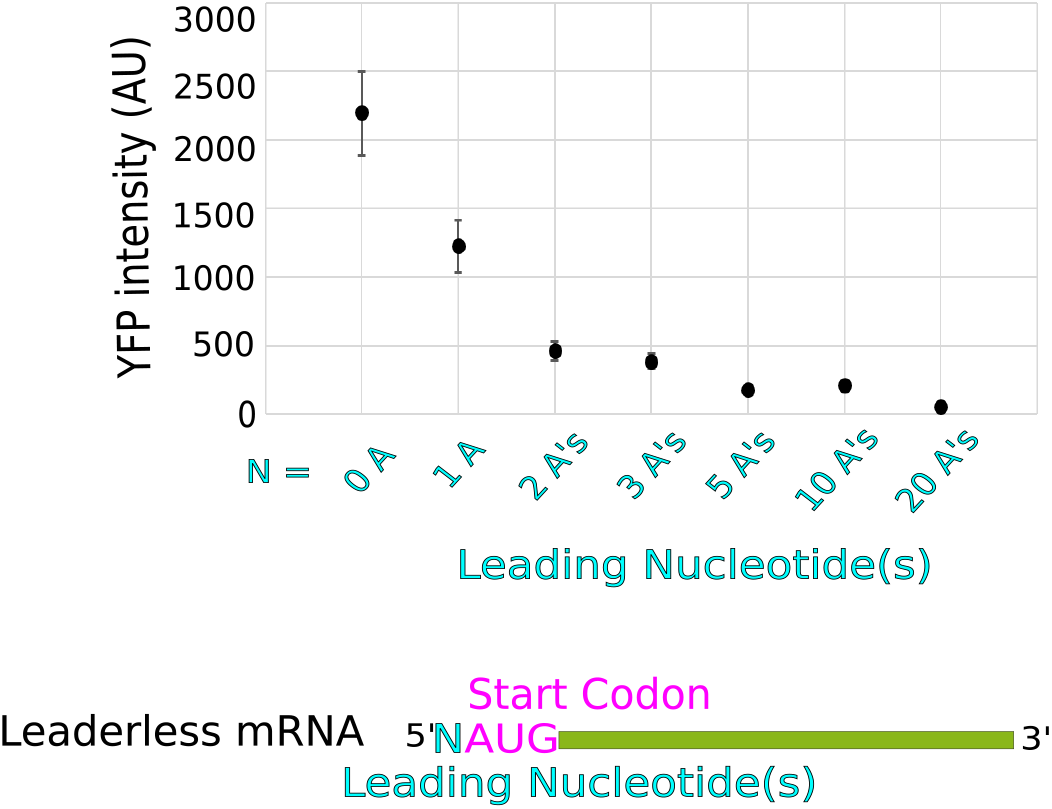
Leaderless mRNAs are inhibited by additional upstream nucleotides. Leaderless mRNA *in vivo* translation reporters were generated with variable number of leading nucleotides on the X-axis and their average YFP intensity per cell were measured (Raw data can be found in Table S1). Error bars represent three biological replicates.

### Leaderless mRNA TIR determinants affect translation efficiency of natural leaderless mRNAs

Because the *in vivo* translation initiation reporters were all synthetic constructs, we explored the extent to which each mRNA feature (ΔG_unfold_, start codon identity, and leader length) occur in natural *C. crescentus* leaderless mRNAs. As noted previously, ΔG_unfold_ is significantly lower for leaderless mRNAs than for other mRNA types (Fig 1B). To analyze the role of start codon selection, we calculated the fraction of AUGs at the 5’ end of all *C. crescentus* leaderless mRNAs and of the random chance of finding each start codon based on the genomes’ GC percentage. This analysis revealed a strong enrichment of AUGs at the 5’ end of *C. crescentus* leaderless mRNAs as compared to random, and a slight enrichment of the GUG near cognate start codons (Fig 5A). While GUG TIR reporters yielded similar TIR reporter levels to UUG and CUG, it is possible that the lack of U and C at the +1 of *C. crescentus* leaderless mRNAs is due to their low abundance in TSSs (40). Of all the leaderless mRNAs, only 4.4% (17/385) are initiating with non-AUG start codons as compared to the leadered mRNAs of which 27.23% (989/3632) of genes initiate with non-AUG start codons (Table S3). Since these near cognate start codons were translated much more poorly than AUG in our translation initiation reporters, it’s possible that for leaderless mRNAs there’s a positive selection for the AUG start codon and a negative selection for near-cognate start codons. Additionally, by exploring the length of mRNAs, we noticed that there was a much greater occurrence of leaderless mRNAs than mRNAs with short leaders <10nt (Fig 5B). Additional leader nucleotides were strongly inhibitory of leaderless translation, and only 8 contain SD motifs, suggesting some of these short-leadered mRNAs may be poorly initiated.

**Figure 5.**
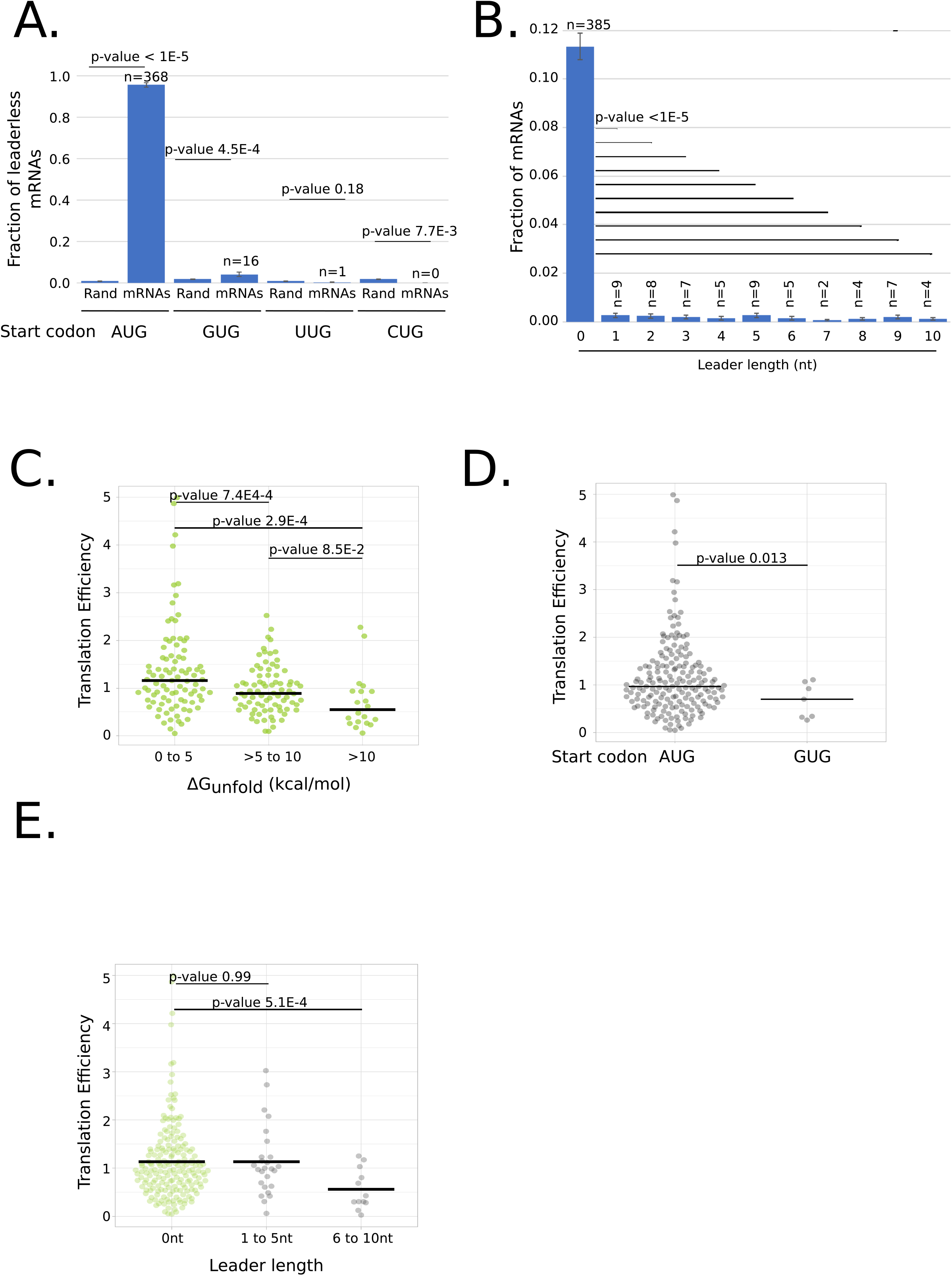
ΔG_unfold_, start codon identity, and leader length correlate with translation efficiency (TE) across native leaderless mRNAs. A.) Bar graph showing the fraction of leaderless mRNAs starting with AUG, GUG, UUG and CUG start codons. Also shown are the random chances of trinucleotides being AUG, GUG, UUG and CUG calculated based on GC content (67%) of *C. crescentus* genome. P-values were calculated based on a two-tailed Z-test. B.) Bar graph showing the fraction of leaderless mRNAs and mRNAs with 5’ untranslated region (UTR) of length 1 to 10 (as determined in (40)). mRNAs containing Shine-Dalagarno sites were excluded from this analysis. P-values were calculated based on a two-tailed Z-test of each leader length compared to leader length 0. C.) Violin plot of translation efficiency (TE) as measured by ribosome profiling(54) of natural leaderless mRNAs binned in three groups depending on ΔG_unfold_ values (0-5, 5-10, and >10 kcal/mol). P-values based on t-test (two tailed, unequal variances). D.) Violin plot of TE as measured by ribosome profiling(54) of natural leaderless mRNAs starting with AUG and GUG. P-values were calculated based on a t-test (2-tailed, unequal variance). E.) Violin plot showing the TE as measured by ribosome profiling(54) on the Y-axis of leaderless mRNAs (green) and with leaders of varying length (1–10) shown in grey. P-values were calculated based on t-test (2-tailed, unequal variance).

To estimate the effects of each mRNA feature (ΔG_unfold_, start codon identity, and leader length) on natural leaderless mRNA translation, we next analyzed ribosome profiling data of the *C. crescentus* mRNAs (27). Here, we utilized translation efficiency measurements which approximate the relative number of ribosome footprints to mRNA fragments from the same cell samples (54). In total, translation efficiency data for 191 leaderless mRNAs and 38 short leadered mRNAs (1-10 leader length) were obtained for cells grown in PYE media (27). We separated leaderless mRNAs into three groups based upon their ΔG_unfold_ values (0-5, 5-10, and >10 kcal/mol) and compared their translation efficiency. The median translation efficiency was reduced as the ΔG_unfold_ increased (Fig 5C) (median= 1.2 for 0-5 kcal/mol, median= 0.89 for 5-10 kcal/mol, median= 0.54 for >10 kcal/mol), similar to the dependence observed in the synthetic translation reporters (Fig 2B). For start codon identity, we noticed that a majority of leaderless mRNAs with near-cognate start codons had translation efficiencies that were not measurable because their genes contained an additional upstream TSS. However, for the 7 GUG mRNAs whose translation efficiency was measured, the median (0.70) was lower than that of the AUG initiated leaderless mRNAs median (0.97) (Fig 5D), in line with the findings of the synthetic reporters (Fig 3). Finally, we compared the translation efficiency of leaderless mRNAs with those with very short leaders (Fig 5E). Since 8 of these mRNAs with short leaders contain SD sequences in the leader, we removed these RNAs from the analysis because we expect them to initiate translation by the canonical mechanism. As leader length increases, we generally observed that the TE tends to decrease (Fig 5E), again in line with the synthetic reporters (Fig 4). Overall these data suggest that the effects of ΔG_unfold_, start codon identity, and leader length observed in the synthetic translation initiation reporters are also observed across natural *C. crescentus* leaderless mRNAs.

Many RNAs present in the *C. crescentus* transcriptome are not initiated as leaderless mRNAs, so we explored the relative fraction of 5’ AUG trinucleotides in all classes of RNAs (Fig 6A). As noted previously, leaderless mRNAs are highly enriched in AUG codons (Fig 5A). Surprisingly, leadered mRNAs contain a similar fraction of 5’ AUGs as would be predicted from the genome’s GC%, which is also observed in small non-coding RNAs (sRNAs), and anti-sense RNAs (asRNAs). Conversely, tRNAs and rRNAs contain zero cases with a 5’ AUG. To explore why these RNAs are not initiated as leaderless mRNAs, we calculated the ΔG_unfold_ of each class of 5’ AUG containing RNA (Fig 6B). If these 5’ AUGs found in non-leaderless RNAs were inaccessible to ribosomes, it would be permissible for this sequence to be present at the 5’ end without causing aberrant initiation. Indeed, for the RNAs with 5’ AUGs, we observe that leaderless mRNAs have a low ΔG_unfold_ (median= 5.0), while leadered mRNAs (median= 9.5), sRNAs (median = 14), and asRNAs (median = 9.0) all contain a significantly higher ΔG_unfold_ values. This suggests that RNAs with inaccessible 5’ AUGs are blocked from leaderless mRNA initiation.

**Figure 6.**
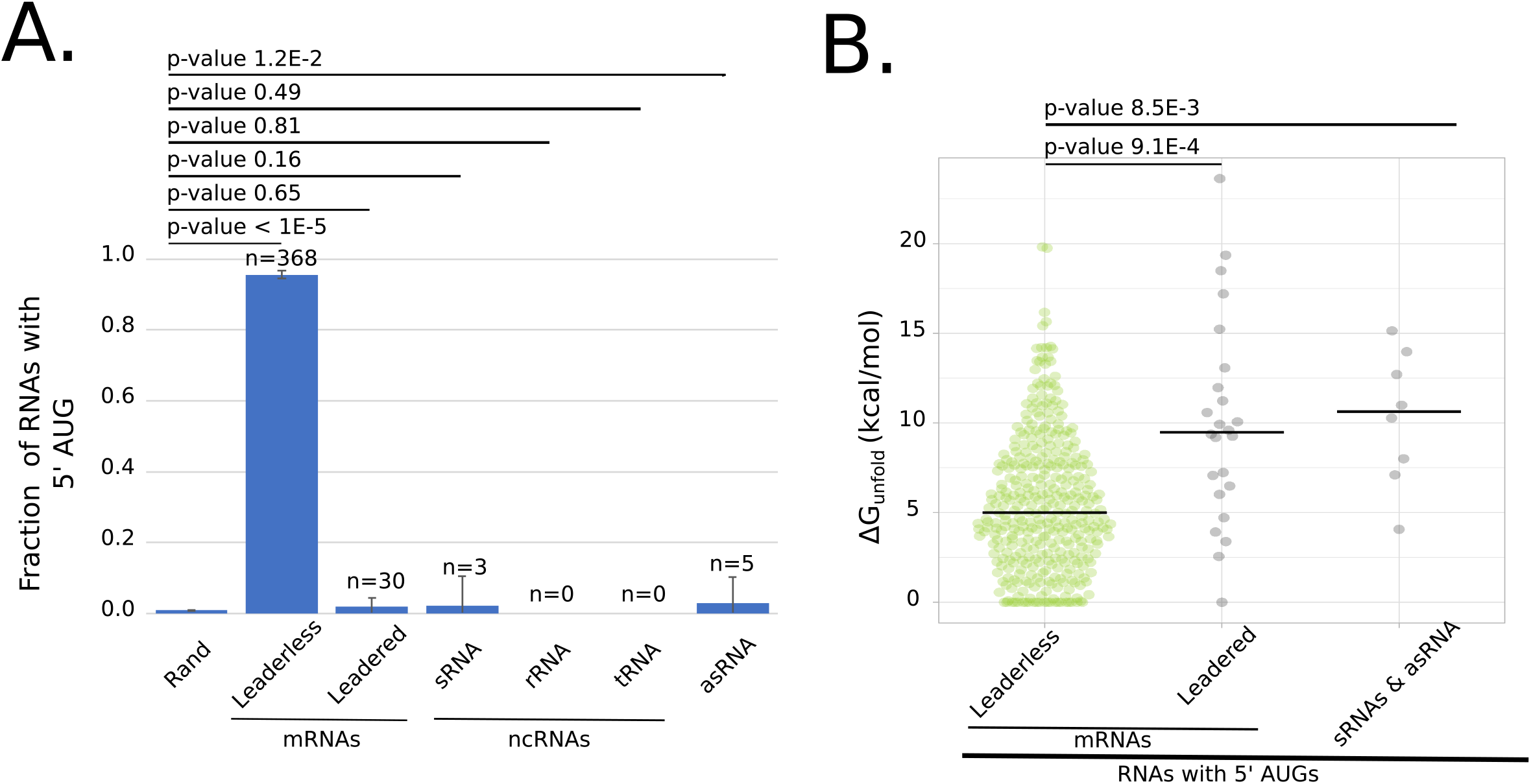
Non-coding RNAs with 5’ AUGs are rare and have higher ΔG_unfold_. A.) Bar graph showing the fraction of natural leaderless mRNAs starting with trinucleotide AUG and other types of RNAs starting with trinucleotide AUG, but not initiated at that AUG (leadered mRNAs, sRNAs, rRNAs, tRNAs and asRNAs). Also shown is the random chance of trinucleotide being AUG out of 10000 nucleotides; calculated based on GC content of *C. crescentus* genome. P-values were calculated using a two-tailed Z-test with each RNA class compared to the random probability of 5’ AUG. B.) Violin plot showing ΔG_unfold_ of natural leaderless mRNAs starting with AUG (green) and other types of RNAs starting with AUG, but not initiated at that AUG (leadered mRNAs, RNAs and asRNAs) (shown in grey). P-values were calculated based on a T-test (2-tailed, unequal variance).

**Figure 7.**
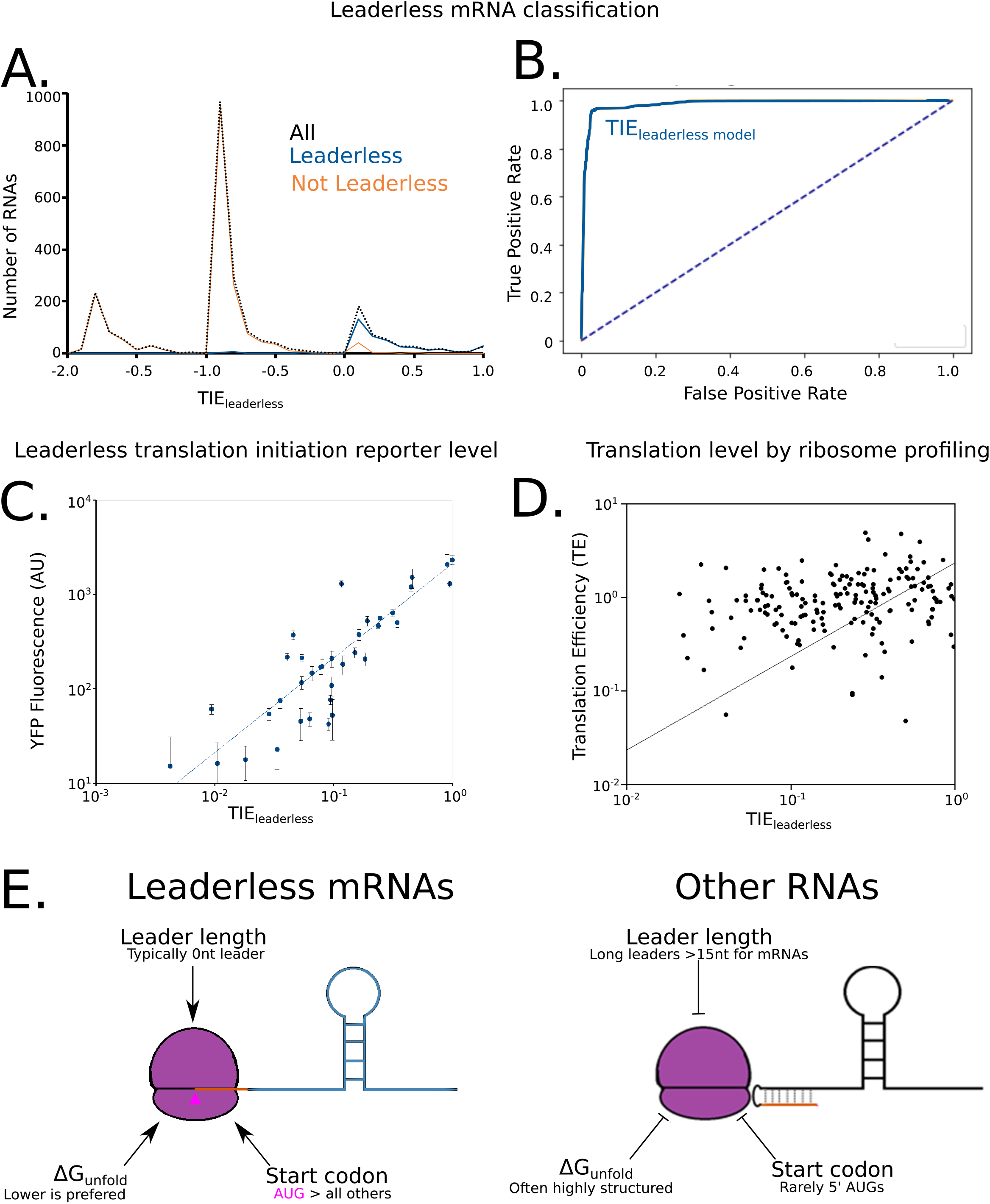
A combinatorial model accurately predicts translation of leaderless mRNAs. A.) Line graph showing the predicted TIE_leaderless_ scores on the X-axis and the number of RNAs on the Y axis. The solid blue line represents natural leaderless mRNAs. The orange line represents the RNAs that are not leaderless RNAs. The black dotted line represents all RNAs. RNAs with short leaders are shown in Fig S2. B.) ROC curve (shown in solid blue, with “random” shown as a dotted line) with true positive rate on Y-axis and false positive rate on X-axis. The area under curve (A.U.C.) was calculated to be 0.99 for classification based on the TIE_leaderless_ score and 0.68 for classification based solely on presence of a 5’ AUG (Fig S3). C.) TIE reporter levels compared to TIE_leaderless_ scores. For the leaderless TIE reporters tested (Table S1) the YFP reporter level (Y-axis) is plotted compared to the TIE_leaderless_ (X-axis). The trendline is the result of a least-squares fit yielding a slope of 2050 A.U. with R^2^=0.87. Error bars represent the standard deviation of at least three biological replicates. D.) Translation efficiency (TE) of leaderless mRNAs (Y-axis) is plotted compared to TIE_leaderless_ (X-axis). The trendline is the result of a least-squares fit yielding a slope of 0.71 and R^2^=0.06. E.) Model design showing ribosome binding to the AUG trinucleotide (pink triangle) at the 5’ end when it is highly accessible as shown in the left. The ribosome binding is prevented when the region becomes more structured and the accessibility decreases.

Due to the strong involvement of ΔG_unfold_, start codon identity, and leader length in *C. crescentus* leaderless mRNA TIRs, we also explored these features across organisms. As already shown in figure 1C, the ΔG_unfold_ for leaderless mRNAs is markedly lower than leadered mRNAs in all species analyzed with the exception of *M. musculus* mitochondria. The low ΔG_unfold_ in the mitochondria may be due to alterations in the translation initiation mechanism of leadered mRNAs by their highly proteinaceous ribosomes (55). Across these organisms, ribosome profiling and total-RNA-seq performed in *M. smegmatis* (16) and *H. volcanii* (44), allowing the comparison of how ΔG_unfold_ tracks with translation efficiency. As observed for *C. crescentus*, as ΔG_unfold_ increases, a drop in the TE in leaderless mRNAs in both *M. smegmatis* and *H. volcanii* (Fig S4). We hypothesize that the low overall ΔG_unfold_ observed for leaderless mRNAs across all species suggests that ribosome accessibility is a key feature for leaderless mRNAs across organisms. In *M. smegmatis* and *H. volcanii* the observed drop in TE from the 0-5 bin and 5-10 bins were smaller than observed for *C. crescentus*, which may be due to the higher growth temperatures of these organisms. Next, we explored the distributions of leader lengths across species (Table S4). As observed in *C. crescentus*,mRNAs with short leaders have a highly skewed distribution across species, with leaderless mRNAs showing the largest peak, with a small fraction of mRNAs containing a 1nt leader, and a much smaller population of mRNAs observed with a ≥2nt leader (Table S4). The lower abundance of mRNAs with very short leaders is likely to be due to their poorer translation levels observed across organisms (FigS4) as this has even been observed with *E. coli* leaderless TIR reporters (37). To explore this possibility, we compared TE across organisms. As observed in *C. crescentus, M. smegmatis* and *H. volcanii* TE is also markedly decreased in mRNAs containing short leaders (Fig S4). While *C. crescentus* leaderless mRNA TE was not significantly lower than the 1-5nt bin, both *M. smegmatis* and *H. volcanii* showed sharper drops in TE. While *C. crescentus* mRNAs with 6-10nt leaders showed a significant drop in TE, neither *M. smegmatis* or *H. volcanii* showed a significant decrease. This discrepancy may be explained by a low sample size in *M. smegmatis*, where only 3 mRNAs were identified in the 5-10nt bin, while in *H. volcanii* the 5-10nt bin distribution contained a single outliner whose TE was measured to be >30. Finally, we examined start codon identities for leaderless mRNAs across species (Table S5). Here only *H. volcanii* was similar to *C. crescentus* with a strong bias in the AUG start codon for leaderless mRNAs (Table S5). *M. tuberculosis* and *M. smegmatis* both contained GUG start codons in leaderless mRNAs with similar abundance to AUG (Table S5). In the *M. musculus* mitochondria, AUG is the most common start codon across mRNAs, however, AUG is less common in leaderless mRNAs, and the summed use of GUG, AUC, AUU, and AUA makes near-cognate start codons more abundant than AUG initiated leaderless mRNAs (Table S5). Interestingly, mitochondrial RNAP has been found to initiate transcription efficiently with NAD+ and NADH (56), which has the potential to alter start codon selection by the translation machinery. As observed in *C. crescentus, M. smegmatis* and *H. volcanii* both showed a lower TE for leaderless mRNAs starting with GUG as compared to AUG (Fig S4). The magnitude of the reduced TE for GUG initiated leaderless mRNAs is significantly smaller in *M. smegmatis* (1.1 AUG, 0.91 GUG) as compared to *H. volcanii* (2.5 AUG, 0.82 GUG), which is in line with previous data showing that GUG initiates with similar efficiency to AUG in leaderless mRNAs in Mycobacteria (16). Overall, these data suggest that the effects of ΔG_unfold_, start codon identity, and leader length have similar effects on leaderless mRNA translation across species. However, minor idiosyncratic differences in the frequency and magnitude of each leaderless mRNA feature on TE were observed across species, likely arising from differences in the translation machinery.

### Three component model describes leaderless mRNA start codon selection

In order to understand how the mRNA determinants combine to dictate leaderless mRNA translation, we built a computational model based upon the three features (ΔG_unfold_, start codon identity, and leader length) and explored its ability to describe leaderless mRNA start codon selection and efficiency of leaderless mRNA translation initiation. From our synthetic *in vivo* translation initiation reporters, we performed curve fitting to assess the relative effect of each feature on TIE. For each feature (ΔG_unfold_, start codon identity, and leader length) the highest reporter level measured in each mutant set was normalized to 1 before curve fitting. ΔG_unfold_ data was fit to an exponential equation (TIE_ΔGunfold_ = e^(-t*0.354)^) where t is ΔG_unfold_(kcal/mol), R^2^ = 0.78), leader length data was fit to a power equation (TIE_leader length_ = 0.45×(i^-0.92^) where i is leader length >0, R^2^ = 0.92, and TIE_leader length_=1 for i=0), and TIE_start codon_ was based directly on reporter levels for each near-cognate start codon (Fig 3) and all other codons were given a value of 0 (Table S1). For each mRNA feature, we therefore generated a function that could calculate the relative TIE of any RNA in *C. crescentus* based upon the mRNA sequence. We then built a computational model in which the three features were assumed to be independent from each other to calculate a summed TIE. In this model, we set the maximum TIE to 1, and then subtracted the effects of the sequence feature as measured from the *in vivo* translation reporters in equation 2 (TIE_Leaderless mRNA*(k)*_ = Max TIE (1) - (1-TIE_ΔGunfold_) - (1-TIE_start codon identity(j)_) − (1-TIE_leader length(i)_) where k= a given RNA transcript, j=start codon identity, and i=leader length(nt). Using equation 2 we predicted the TIE for each RNA in the *C. crescentus* transcriptome (Fig 7A). For all RNAs, we successively scanned for the closest AUG or near cognate start codon to the 5’ end and used this for the TIE calculation. RNAs known to be initiated as leaderless mRNAs (27,40) yielded higher TIE scores (median = 0.15, σ = 0.35), while TIE scores for all other RNAs were typically lower (median = −0.95, σ = 0.45). To estimate the utility of this model at classifying leaderless mRNAs, we used a ROC analysis (Fig 7B). The ROC area under the curve for the TIE_leaderless_ model was equal to 0.99, which significantly outperforms identifying RNAs with 5’ AUGs (ROC A.U.C. 0.68) suggesting the TIE_leaderless_ model can accurately classify those RNAs that are initiated as leaderless mRNAs with high accuracy and precision. The success of this simple TIE_leaderless_ model to classify leaderless mRNAs based on the combinations of ΔG_unfold_, start codon identity, and leader length suggests that these mRNA features combinatorically control translation initiation on leaderless mRNAs.

In addition to the classification of RNAs as leaderless mRNAs we also explored how well the TIE_leaderless_ model predicted translation initiation efficiency. Here, the translation initiation reporters generated were all scored with the TIE_leaderless_ model and compared to their YFP fluorescence. Since TIE_leaderless_ scores below zero are not physically possible, those with negative TIE_leaderless_ values were set to zero to signify they are not predicted to be translated. As expected, the TIE_leaderless_ score correlates strongly to the YFP reporter levels (R^2^=0.87) with a slope of 2050 A.U. We then compared the TIE_leaderless_ scores to the TE as measured by ribosome profiling of the natural leaderless mRNAs. Since natural leaderless mRNAs encode many genes with diverse codon usages, a poorer correlation was obtained with TE (R^2^=0.06, slope=0.71 A.U.) than with the TIE reporters (Fig 7D). The correlation of the TIE_leaderless_ model at predicting ribosome profiling TE (R=0.25) is the same as observed for the RBS calculator model of initiation and *E. coli* ribosome profiling data (R=0.25) (57). Since the TIE reporters all code for YFP with near-identical codon usage, and the natural mRNAs have variable codon usage frequencies, it is possible that translation elongation differences between natural ORFs also impact translation efficiency. Indeed, translation elongation rates have been estimated to be rate limiting *in vivo* in other bacteria (58,59). In addition, ribosome occupancy of stalled ribosomes can complicate the analysis of ribosome profiling data, making the interpretation rather difficult. While it is objectively harder to quantitatively predict translation levels, the TIE_leaderless_ model performs rather well.

## DISCUSSION

Here we provide the first systematic analysis of mRNA structure content, start codon identity, and leader length on the initiation of leaderless mRNAs (Fig 7E). Importantly, this study was performed using the bacterium *C. crescentus* which is adapted to efficient leaderless mRNA initiation (27). As has been observed for leadered mRNAs (19,46), mRNA structure content at the leaderless TIR hinders leaderless mRNA translation initiation, suggesting that ribosome accessibility is a key feature for leaderless mRNAs. As previously observed in *E. coli*, changes in start codon identity from the preferred “AUG” and presence of leader nucleotides leads to a significant reduction of TIE for *C. crescentus* leaderless mRNAs. Using these quantitative data, we generated a combinatorial TIE_leaderless_ model that predicts the ability of an RNA to initiate as a leaderless mRNA from the individual effects of these features which can be computed for any RNA in the transcriptome. This TIE_leaderless_ model both accurately and sensitively predicts the ability of all RNAs in the *C. crescentus* transcriptome to initiate as leaderless mRNAs. While a 5’ AUG is highly enriched in leaderless mRNAs and only rarely observed in non-coding RNAs (Fig 6A), non-coding RNAs containing 5’ AUGs utilize a high ΔG_unfold_ to prevent aberrant translation initiation (Fig 6B). Additionally, very short leaders which were found to inhibit leaderless mRNA initiation, are selected against in leaderless mRNAs and are common in 5’ regions of non-coding RNAs containing non-initiating AUGs. Finally, leaderless mRNAs are much more selective for AUG start codons than are leadered mRNAs, suggesting that the additional stabilization of the translation initiation complex provided by the SD-aSD base pairing helps facilitate initiation on near-cognate start codons.

Leaderless mRNAs have been found to initiate translation in bacterial, archaeal, and both cytoplasmic and mitochondrial eukaryotic ribosomes (17,28,60) suggesting that leaderless initiation is an ancestral initiation mechanism. It is therefore possible that the TIE_leaderless_ model generated here in *C. crescentus* may also perform well across organisms. Indeed, even a few nucleotides preceding the AUG inhibit leaderless mRNA translation initiation in *C. crescentus, E. coli*, and mammalian mitochondria (37,39). The strong inhibition of leaderless mRNA translation by TIR secondary structure is likely why leaderless mRNAs in mitochondria have been found to lack 5’ secondary structures (28). *C. crescentus* shares a similar preference for 5’ AUGs to *E. coli* for leaderless mRNA initiation (33). Interestingly, in the *Mycobacteria*, GUG start codons are much more abundant in leaderless mRNAs and tend to be initiated more similarly to AUG codons in this organism (16). *Mycobacterium* GUG initiated leaderless mRNAs tend to code for short regulatory ORFs (16), as opposed to ORFs encoding functional genes in *C. crescentus*. This suggests that there are likely to be some species-specific differences in leaderless mRNA features arising from the differences in the translation initiation machinery. Indeed, across prokaryotes, 79% of predicted leaderless genes contain AUG as the start codon, whereas GUG, UUG and others are found with an average of 10%, 6% and 3% respectively (13). Surprisingly, leaderless mRNAs across organisms appear to initiate with assembled 70S/80S ribosomes (31,61–63), further suggesting a conserved mechanism of initiation. Therefore, an important goal moving forward will be to determine how broadly across organisms this TIE_leaderless_ model might apply. Based upon the observations described here, it is likely that these features (ΔG_unfold_, start codon identity, and leader length) will combine similarly across species to define leaderless mRNA TIRs. However, due to differences in the translation initiation machinery across organisms, the specific hierarchy of mRNA features will need to be experimentally determined for a given species in order to generate a TIE_leaderless_ model that accurately classifies leaderless mRNAs.

## AVAILABILITY

Transcript architecture for the *Caulobacter crescentus* genome (Updated operon map (updated 4/24/2020)) was obtained from biochemicalphysics.com/resources.

## ACCESSION NUMBERS

Not applicable.

## SUPPLEMENTARY DATA

Supplementary Data are available at NAR online.

## ACKNOWLEDGEMENT

We thank members of the Schrader lab for critical feedback.

