## Supplemental Figures and tables for "A combination of mRNA features influence the efficiency of leaderless mRNA translation initiation": Supplement.docx

**Supplementary Figure Legends**

**Figure S1. ΔGunfold analysis using the same ribosome footprint size for leaderless and leadered TIRs.** For the same RNAs in Figure 1B, we calculated the ΔGunfold using the same 13nt ribosome footprint from leaderless mRNAs constrained as single stranded for all leadered mRNAs. This ensures the constrained single-stranded region is of equal size for both leadered and leaderless mRNA TIRs. P-values were calculated using a 2-sided t-test with unequal variance.

**Figure S2. A combinatorial model classifying mRNAs with very short leaders.**

Line graph showing the predicted TIE_leaderless_ mRNA scores on the X-axis and the number of RNAs on the Y-axis. Only mRNAs with a 5′ UTR of <16 nt without an SD sequence since the exact mechanism of translation initiation is not understood for these RNAs.

**Figure S3. R.O.C. A.U.C. analysis of the TIE_leaderless_ model leaderless mRNA classification and a model of leaderless mRNA classification based only on presence of a 5’ AUG.** TIE_leaderless_ model is shown with a solid blue line, 5’ AUG classification model is shown with a solid grey line, and random classification is shown with a dark blue dashed line.

**Figure S4. ΔGunfold, start codon identity, and leader length correlate with translation efficiency (TE) across native leaderless mRNAs in** ***M. smegmatis* and *H. volcanii*.**

Violin plot of translation efficiency (TE) as measured by ribosome profiling of natural leaderless mRNAs in *M. smegmatis* and *H. volcanii* across different bins of ΔG_unfold_ values and start codons (AUG and GUG). Also, translation efficiency (TE) of natural leaderless mRNAs compared with short leaders in these two organisms.

**Figure S5. Transcriptional reporters of +1nt variants.** Transcriptional reporters containing a mutant version of 5’ leader sequence of *Caulobacter* gene CCNA_03971 were generated in the pBXYFPC-2 plasmid. The +1 nucleotide variants were generated by gene synthesis and measured after a 6-hour induction with xylose on an M2G 1.5% agarose pad. At least 100 cells were used for each measurement.

**Supplementary Tables**

**Table S1. TIE_leaderless_ values and reporter measurements for all variants tested and for the *C. crescentus* transcriptome.**

**(XLSX).**

**Table S2. Codon usage frequencies for all the variants in figure 2**

**(XLSX).**

**Table S3. Frequencies of the start codon variants in *Caulobacter crescentus* *(Ccr)* genome**

| Start codon identity | Frequency of start codons in *Ccr* genome | Frequency of start codons in *Ccr* Leadered mRNAs | Frequency of start codons in *Ccr* Leaderless mRNAs |
| --- | --- | --- | --- |
| AUG | 2886/3885 (74.28%) | 2644/3632 (72.80%) | 368/385 (95.58%) |
| GUG | 565/3885 (14.54%) | 555/3632 (15.28%) | 16/385 (4.15%) |
| UUG | 403/3885 (10.37%) | 403/3632 (11.10%) | 1/385 (0.26%) |
| CUG | 26/3885 (0.67%) | 26/3632 (0.71%) | 0/385 (0%) |
| AUC | 3/3885 (0.077%) | 3/3632 (0.082%) | 0/385 (0%) |
| AUU | 1/3885 (0.026%) | 1/3632 (0.027%) | 0/385 (0%) |
| AUA | 1/3885(0.026%) | 1/3632 (0.027%) | 0/385 (0%) |

**Table S4. Distribution of leader lengths in all other organisms analyzed.**

**(XLSX).**

**Table S5. Frequencies of start codon usage in all other organisms analyzed.**

|  | **Frequency of Start Codons** | | | | | | | |
| --- | --- | --- | --- | --- | --- | --- | --- | --- |
| **Start codon identity** | ***Mtb* all mRNAs** | ***Mtb* Leaderless mRNAs** | ***Msm* all mRNAs** | ***Msm* Leaderless mRNAs** | ***Hvo* all mRNAs** | ***Hvo* Leaderless mRNAs** | ***Mmu* mitochondria all mRNAs** | ***Mmu mitochondria* Leaderless mRNAs** |
| AUG | 2542/4264 (59.62%) | 392/777 (50.45%) | 4030/7109 (56.69%) | 409/946 (43.23%) | 2575/2990 (86.12%) | 913/1012 (90.22%) | 9/13 (69.23%) | 3/7 (42.86%) |
| GUG | 1551/4264 (36.37%) | 385/777 (49.55%) | 2807/7109 (39.49%) | 536/946 (56.66%) | 402/2990 (13.44%) | 99/1012 (9.78%) | 1/13 (7.69%) | 1/7 (14.29%) |
| UUG | 168/4264 (3.94%) | 0/777 (0%) | 272/7109 (3.83%) | 1/946 (<0.01%) | 5/2990 (0.17%) | 0/1012 (0%) | 0/13 (0%) | 0/7 (0%) |
| CUG | 0/4264 (0%) | 0/777 (0%) | 0/7109 (0%) | 0/946 (0%) | 3/2990 (0.10%) | 0/1012 (0%) | 0/13 (0%) | 0/7 (0%) |
| AUC | 2/4264 (<0.01%) | 0/777 (0%) | 0/7109 (0%) | 0/946 (0%) | 2/2990 (0.07%) | 0/1012 (0%) | 1/13 (7.69%) | 1/7 (14.29%) |
| AUU | 0/4264 (0%) | 0/777 (0%) | 0/7109 (0%) | 0/946 (0%) | 1/2990 (0.03%) | 0/1012 (0%) | 1/13 (7.69%) | 1/7 (14.29%) |
| AUA | 1/4264 (<0.01%) | 0/777 (0%) | 0/7109 (0%) | 0/946 (0%) | 2/2990 (0.07%) | 0/1012 (0%) | 1/13 (7.69%) | 1/7 (14.29%) |

**Table S6. List of oligos, plasmids, and strains.**

| Strain name | Reverse primer name | Oligo sequence | length | Plasmid |
| --- | --- | --- | --- | --- |
| JS320 | Leaderless_1_mut1 | gctgatgaaatttctttcattcggcacttggtagcgctaacatg | 44 | pBXYFPC-2_Leaderless_1_mut1 |
| JS321 | Leaderless_1_mut2 | gcgctggagatttccttcattcggcacttggtagcgctaacatg | 44 | pBXYFPC-2_Leaderless_1_mut2 |
| JS322 | Leaderless_1_mut3 | gcgctagagatttccttcattcggcacttggtagcgctaacatg | 44 | pBXYFPC-2_Leaderless_1_mut3 |
| JS458 | Leaderless_2_mut1 | gctgatgggcccgttaacagcgggcccattcggcacttggtagcgctaacatg | 53 | pBXYFPC-2_Leaderless_2_mut1 |
| JS459 | Leaderless_2_mut2 | gctgatgggcccgttaacagggggcccattcggcacttggtagcgctaacatg | 53 | pBXYFPC-2_Leaderless_2_mut2 |
| JS460 | Leaderless_2_mut3 | gctgatgggccagttaacagagggcccattcggcacttggtagcgctaacatg | 53 | pBXYFPC-2_Leaderless_2_mut3 |
| JS461 | Leaderless_2_mut4 | gcacttggtccagttaatagtggtcccattcggcacttggtagcgctaacatg | 53 | pBXYFPC-2_Leaderless_2_mut4 |
| JS462 | Leaderless_2_mut5 | gcacttggtcctgttaatagtggtcccattcggcacttggtagcgctaacatg | 53 | pBXYFPC-2_Leaderless_2_mut5 |
| JS463 | Leaderless_3_mut1 | gctgatgtcgtcgttaacagcgacgacattcggcacttggtagcgctaacatg | 53 | pBXYFPC-2_Leaderless_3_mut1 |
| JS464 | Leaderless_3_mut2 | gctgatgtcgtcgttaacagggacgacattcggcacttggtagcgctaacatg | 53 | pBXYFPC-2_Leaderless_3_mut2 |
| JS465 | Leaderless_3_mut3 | gcacttgtcgtcgttaacagcgaggacattcggcacttggtagcgctaacatg | 53 | pBXYFPC-2_Leaderless_3_mut3 |
| JS466 | Leaderless_3_mut4 | gctgatgtcgtcgttaacagcgaactcattcggcacttggtagcgctaacatg | 53 | pBXYFPC-2_Leaderless_3_mut4 |
| JS467 | Leaderless_3_mut5 | gcacttgtagtagttaatagagaagacattcggcacttggtagcgctaacatg | 53 | pBXYFPC-2_Leaderless_3_mut5 |
| JS434 | Leaderless_4_mut1 | gctgatgggcccggcttgcccattcggcacttggtagcgctaacatg | 47 | pBXYFPC-2_Leaderless_4_mut1 |
| JS435 | Leaderless_4_mut2 | gctgatggacccggcttgcccattcggcacttggtagcgctaacatg | 47 | pBXYFPC-2_Leaderless_4_mut2 |
| JS436 | Leaderless_4_mut3 | gcacttggtcccggcttccccattcggcacttggtagcgctaacatg | 47 | pBXYFPC-2_Leaderless_4_mut3 |
| JS437 | Leaderless_5_mut1 | gctgatggcgccggcttcgccattcggcacttggtagcgctaacatg | 47 | pBXYFPC-2_Leaderless_5_mut1 |
| JS438 | Leaderless_5_mut2 | gcacttggcgccggcttagccattcggcacttggtagcgctaacatg | 47 | pBXYFPC-2_Leaderless_5_mut2 |
| JS439 | Leaderless_5_mut3 | gcacttggtgcaggtttagccattcggcacttggtagcgctaacatg | 47 | pBXYFPC-2_Leaderless_5_mut3 |
| JS442 | Leaderless_6_mut1 | gctgatgtcgtttccgacattcggcacttggtagcgctaacatg | 44 | pBXYFPC-2_Leaderless_6_mut1 |
| JS443 | Leaderless_6_mut2 | gcactagtcgtttccgacattcggcacttggtagcgctaacatg | 44 | pBXYFPC-2_Leaderless_6_mut2 |
| JS444 | Leaderless_6_mut3 | gctgatgtcgtttcggacattcggcacttggtagcgctaacatg | 44 | pBXYFPC-2_Leaderless_6_mut3 |
| JS445 | Leaderless_6_mut4 | gctgaagtagtttctgacattcggcacttggtagcgctaacatg | 44 | pBXYFPC-2_Leaderless_6_mut4 |
| JS446 | Leaderless_7_mut1 | gctgatgcggtttcccgcattcggcacttggtagcgctaacatg | 44 | pBXYFPC-2_Leaderless_7_mut1 |
| JS447 | Leaderless_7_mut2 | gctgatgcggtttctcgcattcggcacttggtagcgctaacatg | 44 | pBXYFPC-2_Leaderless_7_mut2 |
| JS448 | Leaderless_7_mut3 | gcacttgccgtttctcgcattcggcacttggtagcgctaacatg | 44 | pBXYFPC-2_Leaderless_7_mut3 |
| JS523 | Leaderless_3_mut5_GUG | gcacttgtagtagttaatagagaagacactcggcacttggtagcgctaacatg | 53 | pBXYFPC-2_Leaderless_3_mut5_GUG |
| JS524 | Leaderless_3_mut5_UUG | gcacttgtagtagttaatagagaagacaatcggcacttggtagcgctaacatg | 53 | pBXYFPC-2_Leaderless_3_mut5_UUG |
| JS525 | Leaderless_3_mut5_CUG | gcacttgtagtagttaatagagaagacagtcggcacttggtagcgctaacatg | 53 | pBXYFPC-2_Leaderless_3_mut5_CUG |
| JS526 | Leaderless_3_mut5_AUC | gcacttgtagtagttaatagagaagagattcggcacttggtagcgctaacatg | 53 | pBXYFPC-2_Leaderless_3_mut5_AUC |
| JS527 | Leaderless_3_mut5_AUA | gcacttgtagtagttaatagagaagatattcggcacttggtagcgctaacatg | 53 | pBXYFPC-2_Leaderless_3_mut5_AUA |
| JS528 | Leaderless_3_mut5_AUU | gcacttgtagtagttaatagagaagaaattcggcacttggtagcgctaacatg | 53 | pBXYFPC-2_Leaderless_3_mut5_AUU |
| JS529 | Leaderless_3_mut5_GGG | gcacttgtagtagttaatagagaagaccctcggcacttggtagcgctaacatg | 53 | pBXYFPC-2_Leaderless_3_mut5_GGG |
| JS530 | Leaderless_3_mut5_1A | gcacttgtagtagttaatagagaagacatttcggcacttggtagcgctaacatg | 54 | pBXYFPC-2_Leaderless_3_mut5_1A |
| JS531 | Leaderless_3_mut5_2A | gcacttgtagtagttaatagagaagacattttcggcacttggtagcgctaacatg | 55 | pBXYFPC-2_Leaderless_3_mut5_2A |
| JS532 | Leaderless_3_mut5_3A | gcacttgtagtagttaatagagaagacatttttcggcacttggtagcgctaacatg | 56 | pBXYFPC-2_Leaderless_3_mut5_3A |
| JS534 | Leaderless_3_mut5_5A | gcacttgtagtagttaatagagaagacatttttttcggcacttggtagcgctaacatg | 58 | pBXYFPC-2_Leaderless_3_mut5_5A |
| JS535 | Leaderless_3_mut5_10A | gcacttgtagtagttaatagagaagacattttttttttttcggcacttggtagcgctaacatg | 63 | pBXYFPC-2_Leaderless_3_mut5_10A |
| JS536 | Leaderless_3_mut5_20A | gcacttgtagtagttaatagagaagacattttttttttttttttttttttcggcacttggtagcgctaacatg | 73 | pBXYFPC-2_Leaderless_3_mut5_20A |
