## Supplementary figures and images for "A combination of mRNA features influence the efficiency of leaderless mRNA translation initiation"

### Fig_S1.pdf

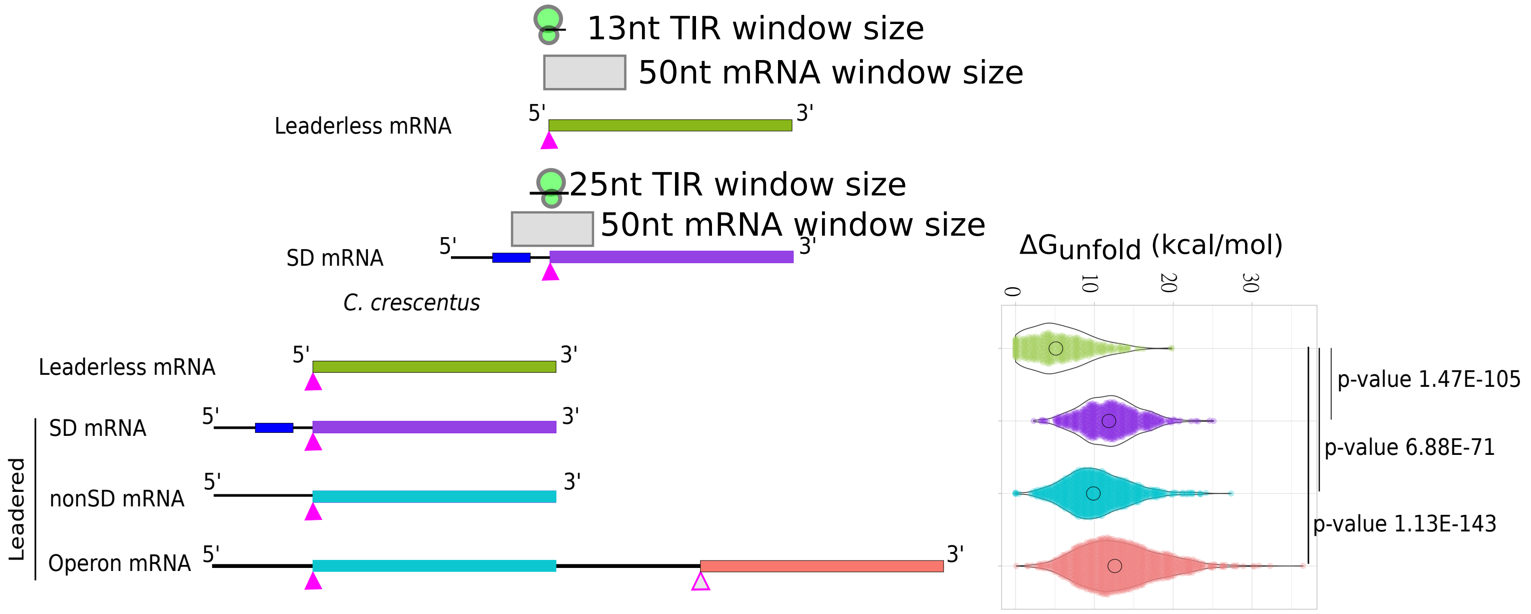

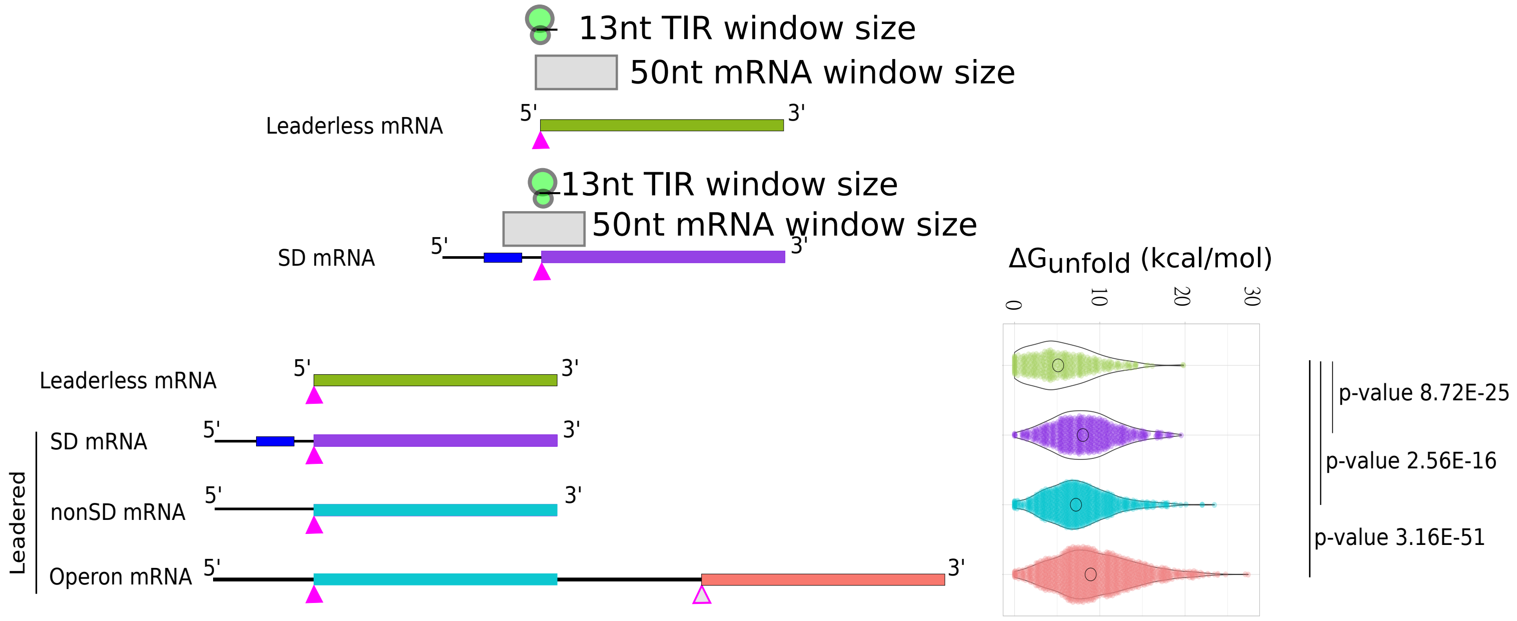

### Fig_S2.pdf

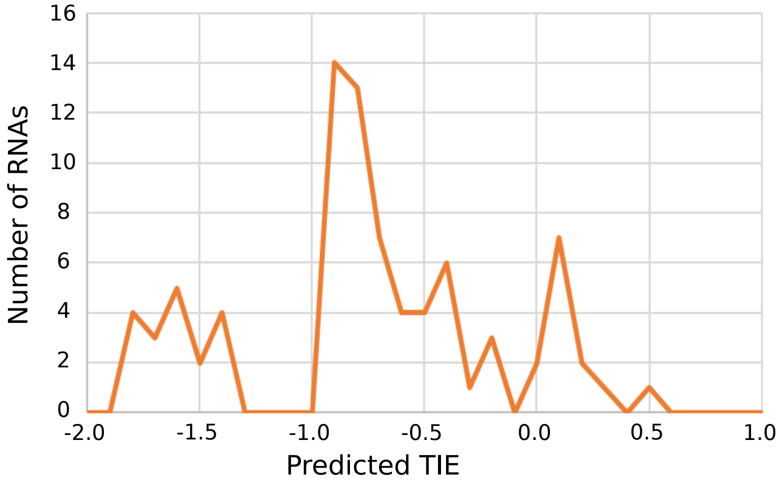

### Fig_S3.pdf

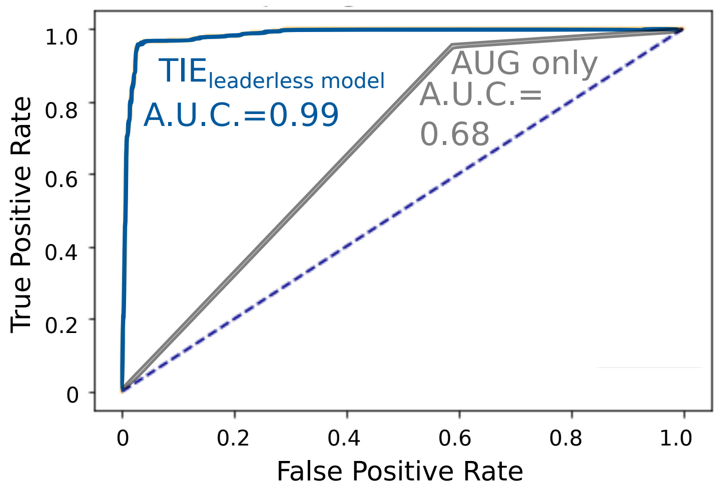

### Fig_S4.pdf

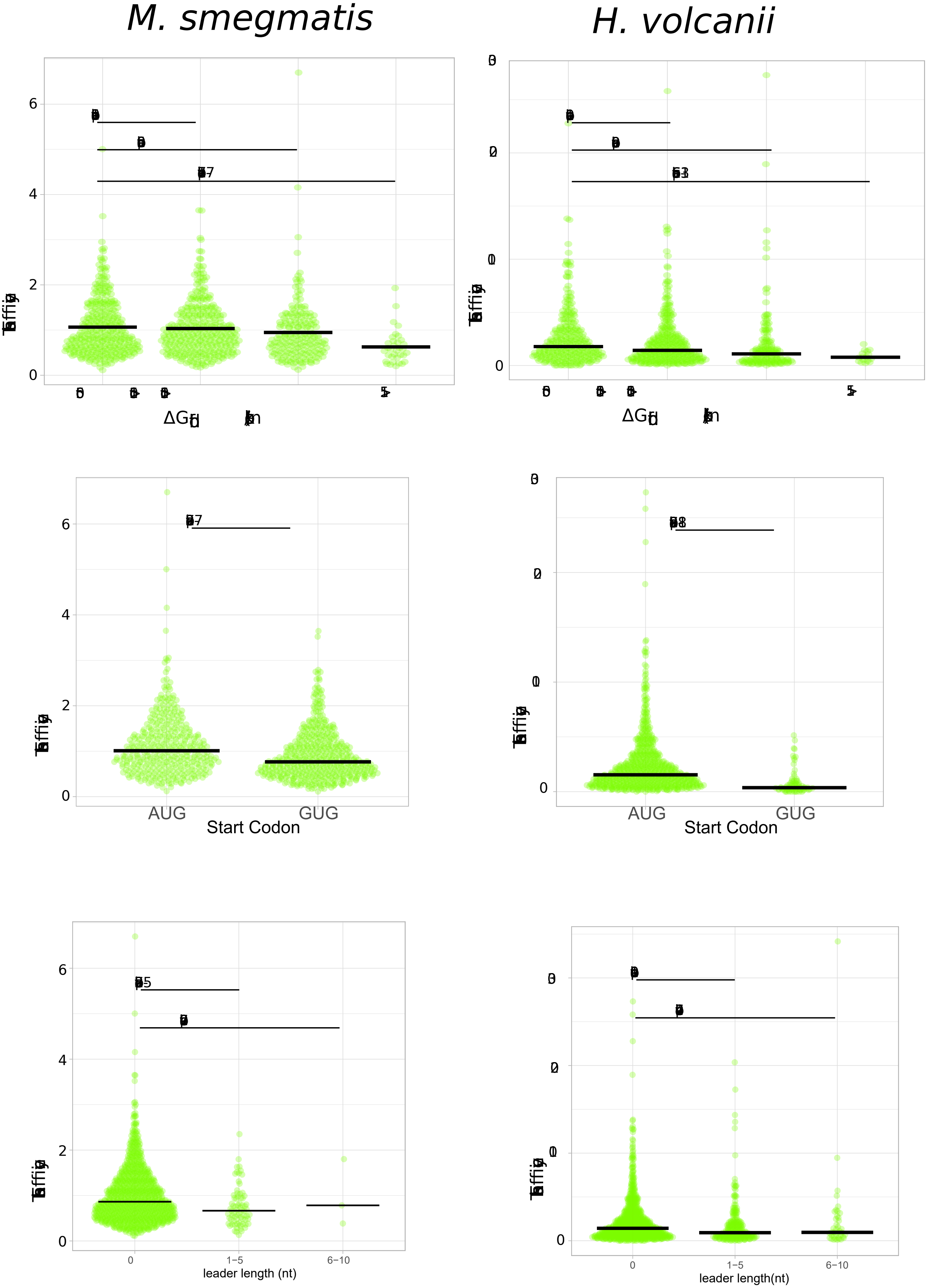

### Fig_S5.pdf

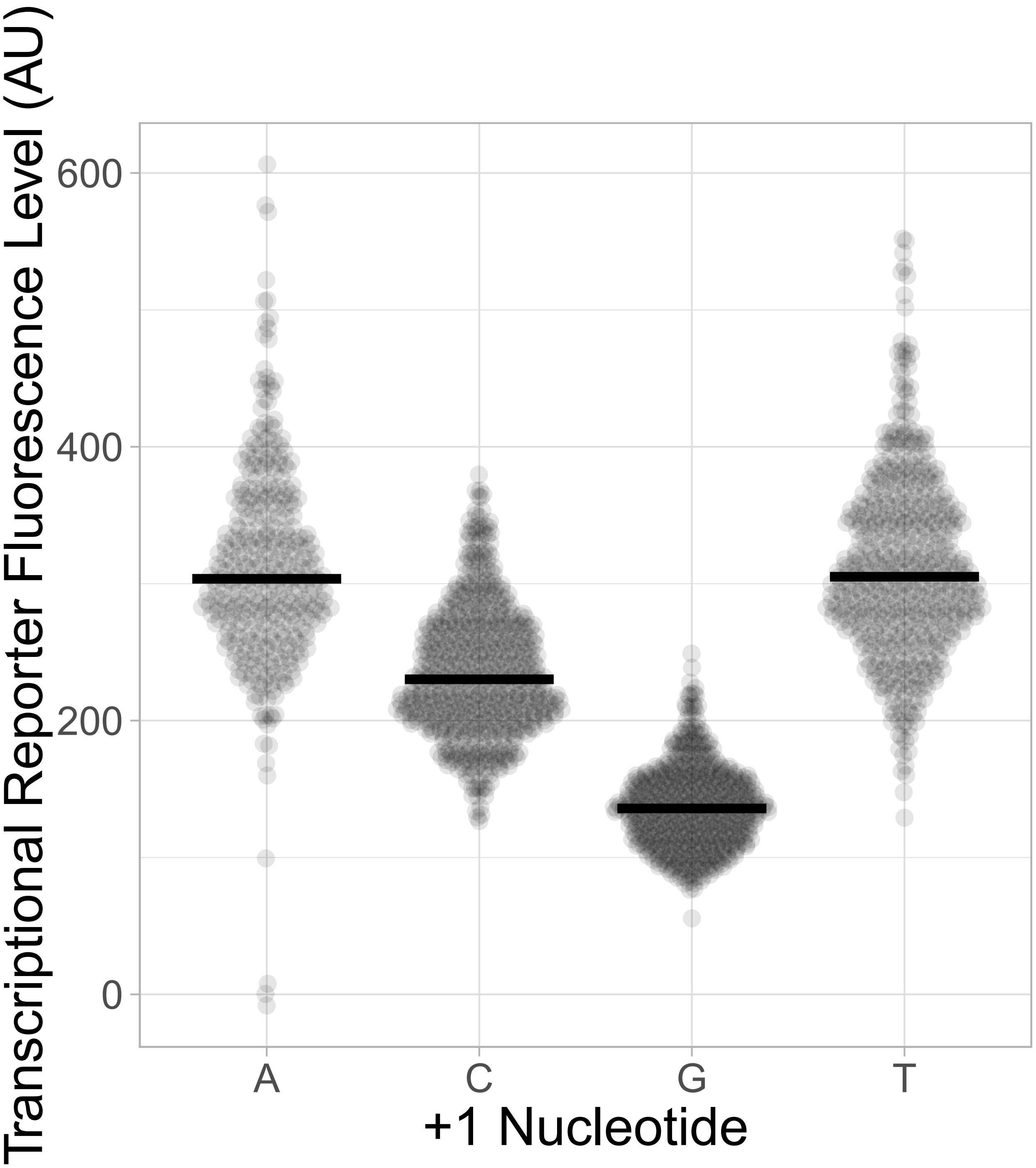
